# Variety is the spice of life: nongenetic variation in life histories influences population growth and evolvability

**DOI:** 10.1101/2024.03.02.583100

**Authors:** Amy B. Forsythe, Sarah P. Otto, William A. Nelson, Troy Day

## Abstract

Individual vital rates are key determinants of lifetime reproductive success, and variability in these rates shapes population dynamics. Previous studies have found that this vital rate hetero-geneity can influence demographic properties including population growth rates, however, the explicit effects of the amount of variation within and the covariance between vital rates that can also vary throughout the lifespan on population growth remains unknown. Here, we explore the analytical consequences of nongenetic heterogeneity on long-term population growth rates and rates of evolution by modifying traditional age-structured population projection matrices to incorporate variation among individual vital rates. The model allows vital rates to be permanent throughout life (“fixed condition”) or to change over the lifespan (“dynamic condition”). We reduce the complexity associated with adding individual heterogeneity to age-structured models through a novel application of matrix collapsing (“phenotypic collapsing”), showing how to collapse in a manner that preserves the asymptotic and transient dynamics of the original matrix. The main conclusion is that nongenetic individual heterogeneity can strongly impact the longterm growth rate and rates of evolution. The magnitude and sign of this impact depends heavily on how the heterogeneity covaries across the lifespan of an organism. Our results emphasize that nongenetic variation cannot simply be viewed as random noise, but rather that it has consistent, predictable effects on fitness and evolvability.

## Introduction

Just as no two snowflakes are alike, neither are two individuals. Ecologists and evolutionary biologists have long sought to capture this among-individual variation in life histories because schedules of individual mortality and reproduction are key determinants of lifetime reproductive success at the individual level and fitness at the population level. Accumulating research shows that this variability among individual mortality and birth rates (collectively referred to as “vital rates”) exists in nearly all populations, from unicellular organisms to mammals (e.g., Armstrong et al. 2018; Beckage and Clark 2003; Hartemink and Caswell 2018; Jouvet et al. 2018; Rotella 2023). There are countless mechanisms that may generate and maintain variability in individual vital rates, from conspicuous differences due to age (Oosthuizen et al. 2021; Tompkins and Anderson 2019), developmental stage (Bright Ross et al. 2020; Deere et al. 2017), size (Armstrong et al. 2018; Rees and Ellner 2019), and genotype (Cressler et al. 2017; Jouvet et al. 2018), to less obvious differences such as unequal allocation of parental care (Dudeck et al. 2018; Smiseth et al. 2007), maternal effects (Fox et al. 2006; Hernández et al. 2020; Jones et al. 2005), developmental hetergoeneity (Cooper and Kruuk 2018; Olijnyk and Nelson 2013), and environmental heterogeneity (Bruijning et al. 2019; Dahlgren et al. 2016; Douhard et al. 2013). Given the overwhelming prevalence of individual heterogeneity, studies that consider its effects on demography can provide invaluable insight about populations and their ecological and evolutionary dynamics.

A central goal across studies of individual heterogeneity is to determine how much, and when, variation among individual life histories matters for population dynamics. Consider a population with heterogeneity in one or more vital rates both among and within age classes (fig. 1). The vital rates that an individual expresses at any given time will depend on its age class and on its phenotype within that age. Some phenotypic categories will allow individuals to express stronger vital rates (e.g., higher survival probabilities or birth rates) than others, and these individuals can be thought of as being in higher “condition” (Forsythe et al. 2021; Ronget et al. 2017). As an individual moves from one age to the next, it may keep its current phenotype, or it may switch to another phenotypic category and obtain a higher or lower condition. The degree of persistence of vital rate phenotypes throughout the lifespan then falls along a scale, with phenotypes that are set at birth and permanent throughout life (“fixed condition,” fig. 1A) at one extreme and phenotypes that may change randomly at any time (“uncorrelated condition^1^,” fig. 1C) at the other (see Cam et al. 2016 and Forsythe et al. 2021 for reviews of terminology). In between these two extremes, “dynamic condition” (fig. 1B) allows the phenotype to persist to some extent but not completely over the lifespan.

**Figure 1.**
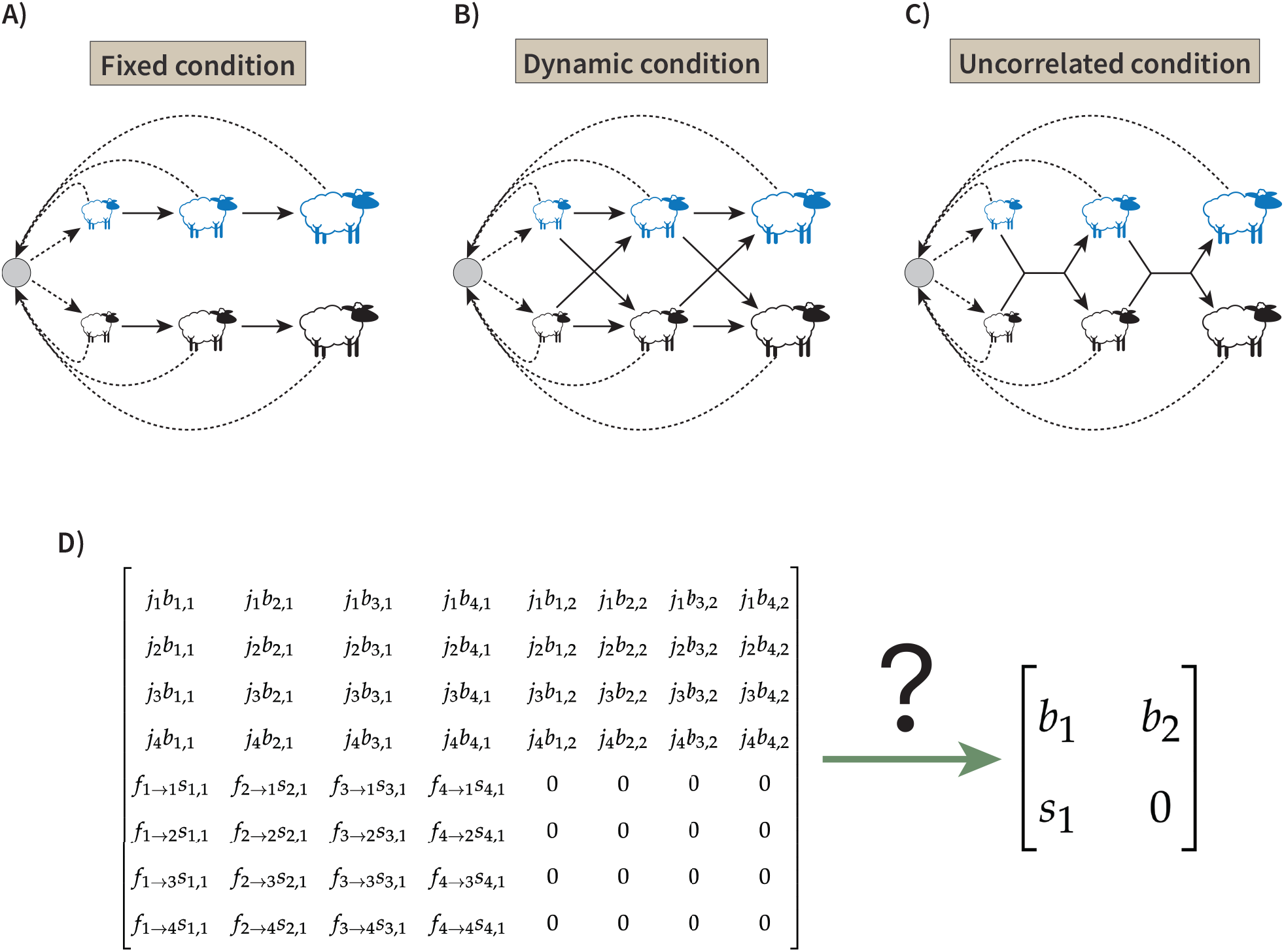
**A-C)** The population model in this study allows individual survival probabilities (solid arrows) and birth rates (dashed arrows) to depend both on an individual’s age class (represented by the individual’s size here) and on its phenotype category (blue or black in this example). Note that all offspring are born into a common “pool” (grey circles) and then assigned a phenotype according to a common newborn probability distribution. This structure generates three potential types of heterogeneity, which are defined by the degree of persistence in individual vital rates throughout the lifespan (focus on the arrows between the sheep): **A**) At one extreme, fixed condition occurs when individual vital rates are set at birth and persist throughout life. For example, seeds that germinate in a particularly good micro-habitat are likely to have higher vital rates than seeds that germinate in poorer environments, and these differences would persist as long as the environmental heterogeneity is maintained. **B)** Dynamic condition allows individuals to transition among phenotypes throughout life, with transition probabilities that can depend on the individual’s current and future age and phenotype. Thus, the phenotype may persist to some extent, but not completely over the lifespan of an individual. For example, heavier animals tend to have higher vital rates (e.g., Bonnet et al. 2017; Hamel et al. 2009; Nussey et al. 2011), however an individual’s body mass may increase or decrease from year to year. **C)** Uncorrelated condition occurs at the extreme of dynamic condition, when an individual’s future phenotype is independent of its current age and phenotype. This type of heterogeneity can result when vital rate phenotypes depend strongly on environmental conditions that vary randomly in a patch from year to year. **D)** The life cycle described in panels A)-C) groups individuals by both age class and phenotypic category, resulting in a large transition matrix (here shown for just two age classes and four phenotypic categories within each age). In this study we develop “phenotypic collapsing” as a method that simplifies this matrix to two age classes and a single, average phenotype in each.

Previous work has shown that these different types of heterogeneity have important impacts on population dynamics (e.g., Acker et al. 2014; Fox 2005; Vattiato et al. 2021; Vindenes et al. 2008, 2012), however a general understanding of their consequences for population growth remains unknown. This is in part because models accounting for heterogeneity become complex at even modest numbers of age and phenotypic categories (fig. 1D), particularly when individual phenotypes can change with age. For example, individuals in higher condition are expected to contribute more to population growth than those in lower condition because of their higher fitness (Coulson et al. 2006; Jansen et al. 2012; Jenouvrier et al. 2015; Zuidema et al. 2009), but these effects last only as long as an individual’s vital rates are maintained. Here we develop a new technique for matrix collapsing (fig. 1D) and use it to gain precise analytical insight for when, and how, different types of vital rate heterogeneity are expected to influence long-term population growth rates (i.e., fitness) and rates of evolution.

In populations structured by age and phenotype (e.g., fig. 1), further complexity arises because there is the potential for both positive and negative covariances in survival and birth rates both across individuals and across ages, which complicates ecological and evolutionary dynamics (Benton et al. 2006; Lee et al. 2017*a*,*b*). Indeed, negative covariances (i.e., “trade-offs”) among vital rates may dampen the effects of individual heterogeneity because they minimize lifetime fitness differences among individuals. These negative covariances are consistent with traditional life-history theory that postulates a fixed energy budget for investment in survival and reproduction, such that an individual that invests heavily in one vital rate will pay a cost with a low value of another (Reznick 1985; Stearns 1989). On the other hand, positive covariances can structure populations into groups of high and low lifetime fitness, where some “lucky” individuals are of high condition because they maintain high values of multiple vital rates throughout life and contribute disproportionately to population growth, compared to individuals in poorer condition (Jansen et al. 2012; van de Pol et al. 2006; Zuidema et al. 2009). Positive covariances are generally expected when there is heterogeneity in resource aquisition among individuals or in resource abundance across environmental patches (Descamps et al. 2016; van de Pol et al. 2006; van Noordwijk and de Jong 1986). While some population models have suggested that covariances among vital rates across the lifespan should have little qualitative effects at the population level unless they are quite strong (González-Suárez et al. 2011; Lee et al. 2017*b*), others have found that both the magnitude and sign of the covariance between survival and birth rates can influence population dynamics (Cressler et al. 2017; Kendall et al. 2011; Stover et al. 2012). In this paper, we show explicitly how the covariance among survival and birth rates over the lifespan of an organism influences long-term population growth rates and rates of evolution.

Previous theoretical studies have considered the effects of individual heterogeneity on long-term population growth, however the results are typically limited either because they rely on numerical analyses, or they assume only one source of heterogeneity (age or phenotype, but not both). Among these, Kendall et al. (2011) developed a population model without age or stage structure in vital rates but allowing two categories of individuals (high and low fixed condition in either survival probabilities or fecundities); heterogeneity in survival probabilities had a “previously unrecognized effect,” increasing the asymptotic population growth rate via “cohort selection,” whereby the higher condition individuals out-survived those in lower condition.

Building on this work, Stover et al. (2012) incorporated fixed condition by setting mortality rates, birth rates, or both to a high or low value in a density-dependent logistic model and found that the low-density growth rate depends on the strength of individual heterogeneity as well as the correlation among vital rates. These models suggest that heterogeneity in vital rates can have a strong influence on population growth when these rates remain constant as an individual ages, however we still lack theory for the influence of such heterogeneity on population growth rates when vital rates can vary with age. This is in part because including heterogeneous vital rates in age-structured populations allows for much more complexity, as the condition of an individual can change over their life span, with the potential for both positive and negative covariances in age-specific survival and fertility rates (fig. 1).

There is a large body of analytical work that explores the effects of individual heterogeneity within age or stage classes on the expectation and variance in the total number of offspring for each newborn individual, i.e., “lifetime reproductive success” (e.g., Jenouvrier et al. 2015; Plard et al. 2012; Snyder and Ellner 2018, 2022; Snyder et al. 2021; Tuljapurkar et al. 2009, 2020; van Daalen and Caswell 2020). Several studies have used these analyses along with measures of generation time to quantitatively estimate how among-individual variation may also influence population growth rates (e.g., Steiner and Tuljapurkar 2012; Steiner et al. 2014). While this question has also been investigated using integral projection models (IPMs), to our knowledge such analyses of population growth using IPMs have remained numerical (e.g., Fung et al. 2022; Kentie et al. 2020; Plard et al. 2016, 2018; Vindenes and Langangen 2015). Here we identify the explicit effects of the amount of variation within and the covariance among survival and birth rates on long-term population growth.

Traditional evolutionary studies often focus on genetic variation that is heritable and so can generate evolutionary change (Huxley 1942; Mayr 1982). Yet, nongenetic factors can be just as, if not more, important than genetics in determining fitness, and these effects of nongenetic variation can scale up to influence ecological and evolutionary dynamics (Cressler et al. 2017; Gomes et al. 2019; Jouvet et al. 2018; Olijnyk and Nelson 2013). For example, Jouvet et al. (2018) identified variation in lifespan and lifetime reproduction within clones of the bacterium *Escherichia coli*, which shifted the clonal population growth rate (Malthusian fitness) away from what would be predicted if all cells had followed the same life-history schedule represented by the mean lifespan and mean lifetime reproduction of the clone. Nongenetic individual heterogeneity also plays a dominant role for determining evolutionary dynamics in *Daphnia pulicaria*, where it has been observed that even in the same environment, there is more variation in mortality, birth, and development rates among individuals within a clonal genotype than there is among genotypes (Cressler et al. 2017; Olijnyk and Nelson 2013). Theory that builds on conventional models to include such nongenetic variation in vital rates can therefore improve our understanding of the ecological and evolutionary dynamics that occur in real populations.

Evolutionary studies of individual heterogeneity within age- and stage-structured populations have compared differences that arise when the heterogeneity is generated by fixed categories (fig. 1A) versus stochastic variability (fig. 1C) among individuals (e.g., Snyder and Ellner 2018; Steiner and Tuljapurkar 2012). However, these stochastic differences are generally assumed to be neutral with respect to selection while the fixed differences are assumed to be genetically based; under this assumption, stochastic variation in vital rates throughout the life course can magnify the effect of random drift in populations and slow the rate of evolution compared to the change expected from fixed genetic differences alone (Coste and Pavard 2020; Steiner and Tuljapurkar 2012; van Daalen and Caswell 2020). Individual-based simulations have suggested a similar effect of nongenetic heterogeneity that persists completely throughout the lifespan, with increasing strengths of nongenetic relative to genetic heterogeneity influencing population growth rates and slowing rates of evolution by natural selection (Cressler et al. 2017). These simulation results depended on the sign of the nongenetic and genetic covariances among vital rates, however the general mathematical relationships between these covariances and rates of evolution and population growth remain unknown. In this study we develop a conceptual unification of the influences of nongenetic heterogeneity on population growth rates and rates of evolution, with a particular emphasis on the impact of the covariance among vital rates throughout the lifespan.

Here, we explore the consequences of different types of nongenetic individual heterogeneity on the long-term growth rate (fitness) of a genotype, allowing us to determine the impact of this heterogeneity on the rate of evolution by natural selection. We modify traditional age-structured population projection matrices (Leslie matrices) to incorporate individual heterogeneity within age classes among survival probabilities and birth rates. We assume that each population projection matrix represents a single, clonal genotype, such that all individual heterogeneity within a matrix is attributed to nongenetic sources. The model includes further biological realism by allowing arbitrary transitions in phenotype over the lifespan. To gain analytical insight we make one key assumption: that the heterogeneity is reset among the offspring in each generation (or, equivalently, at some fixed age), however this could be relaxed in future numerical analyses. This assumption allows us to introduce “phenotypic collapsing” as a method to reduce the complexity associated with adding individual heterogeneity to age-structured models (fig. 1D), while preserving both the transient and asymptotic dynamics of the original system. Genetic variation is then added by considering multiple clonal genotypes, each with their own transition matrix incorporating nongenetic heterogeneity. We compare the long-term growth rate and evolvability across a broad range of genetic and nongenetic covariances and across different strengths of nongenetic variation. The main conclusion is that nongenetic individual heterogeneity strongly impacts the long-term population growth rate and rate of evolution by natural selection. The magnitude and even the sign of this impact depends on whether the heterogeneity is fixed or varying across the lifespan and on how the heterogeneity co-varies across vital rates. Our results emphasize that nongenetic variation cannot simply be viewed as random noise because it has predictable non-neutral effects on fitness and evolvability.

## Methods and Results

We modify classic age-structured population projection matrices to study the evolutionary consequences of nongenetic individual heterogeneity. The population modelled follows the lifecycle in fig. 1, where individuals survive and give birth with probabilities determined by their current phenotype and age. The model assumes that individuals move to the next age class at each time step and may transition to a different phenotype as they age. We assume that the model includes enough age classes that essentially no individuals in the oldest age class survive another time step, an assumption that is required for the phenotypic collapsing method that we develop below.

Our approach begins by considering the fitness of a single genotype with vital rates that are assigned at birth and either permanent throughout life (fixed condition, fig. 1A) or allowed to change throughout the life course (dynamic condition, fig. 1B). Vital rates that vary over the lifespan may depend on the individual’s current age and phenotype or may be be drawn randomly at each age (uncorrelated condition, fig. 1C). We then use this analysis to understand how nongenetic heterogeneity in condition influences the long-term growth rate and rate of evolution by natural selection across multiple genotypes in a population.

### Analysis - Single Genotype

The traditional form of an age-structured population projection matrix assumes that individuals can be classified into states that depend only on their age (Leslie 1945). The full population projection matrix model is needed to predict short-term (“transient”) dynamics, while the long-term (“asymptotic”) dynamics are determined knowing only the largest (“leading”) eigenvalue (*λ*_1_, which gives the asymptotic population growth rate) and its corresponding right and left eigenvectors (which give the stable-age distribution and age-specific reproductive values, respectively) (Caswell 2001). Here we assume that each transition matrix describes an isolated population consisting of a single genotype, and consider *λ*_1_ to be the relevant measure of genotypic fitness. Scaling up to a population of multiple genotypes, the variance among genotypes in *λ*_1_ gives the total genetic variance in fitness (*V*_*G*_), which predicts the rate of fitness increase (“evolvability”) under the assumption of no interaction or mating among genotypes (Fisher 1930).

To explore the fitness consequences of different types of nongenetic individual heterogeneity, we allow both vital rates in the Leslie matrix (survival probabilities, *s*, and birth rates, *b*) to vary among individuals within each age class. Individuals are assigned a value for each vital rate at birth, and these rates are either permanent (fixed condition, fig. 1A) or can vary arbitrarily throughout life (dynamic condition, fig. 1B).

We first develop a simple example for a single genotype with two age classes and two potential values of each vital rate to highlight the structural differences between life histories that are fixed or varying across life and to investigate how these differences influence genotype fitness. We then extend this model to an arbitrary number of age classes and values of each vital rate and describe how the resulting large-scale matrices can be collapsed to matrices with a single set of effective vital rates for each age class, while still providing the complete solution to the original model. Note that this means that the eigenvalues and eigenvectors of the reduced system completely characterize the solution of the original full system (after a maximum of one full lifetime, see Appendix 1). Throughout, we assume that population sizes are large, and so we can ignore stochastic variation in the realized heterogeneity that would be expected in smaller populations (e.g., Gillespie 1973, 1974, 1977). Finally, we conduct a numerical analysis to explore the relative importance of genetic and nongenetic sources of heterogeneity to the mean long-term growth rate and rate of evolution in a population of distinct genotypes.

#### A Simple Example

Consider an age-structured clonal population of a single genotype with no density-dependence and two age classes: non-reproductive juveniles (*J*) and reproductive adults (*A*). The Leslie matrix is 

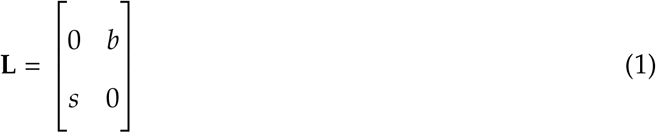

This matrix assumes that the survival rate of juveniles to adults (*s*) and the birth rate of adults (*b*) is the same for all individuals and that all adults die after giving birth.

To incorporate individual heterogeneity in this model, assume that individuals can express one of *ρ* different phenotypes, each with their own unique potential vital rates. Here we use the term “phenotype” to distinguish variants that are produced by nongenetic variation from those that are produced by genetic variation, however the vital rates need not be directly observable. Individuals of phenotype *x* survive with probability *s*_*x*_ and give birth to *b*_*x*_ offspring over a single time step.

For this simple example with two age classes, we consider two phenotypes in the population. The first phenotype is represented by a set of vital rates (*s*_1_, *b*_1_) and the second phenotype by a separate set of vital rates (*s*_2_, *b*_2_). At a given time, individuals can belong to one of four possible states in this model, such that

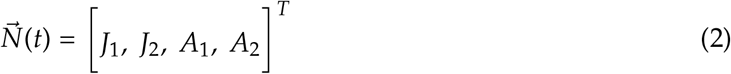

where 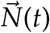 is the state vector at time *t*, and *J*_*x*_ and *A*_*x*_ represent the number of juveniles and adults of phenotype *x* at time *t*, respectively. A core assumption that we make is that the phenotypic heterogeneity is reset in the offspring, with no memory of the parent’s phenotype or age (genetic differences are treated later), so that adults give birth to offspring of phenotype *x* with probability given by *j*_*x*_, regardless of the parent. Further, we let *j*_1_ + *j*_2_ = 1, so that all offspring must be born into one of the two phenotypes and there is no death of newborns from the transition process.

Under fixed condition, an individual’s vital rates are fixed for life (fig. 1A). Thus, individuals may be born from a parent of either phenotype, but once the phenotype is assigned it determines an individual’s survival probability and birth rate for its entire life. The transition matrix for a genotype with fixed condition is therefore 

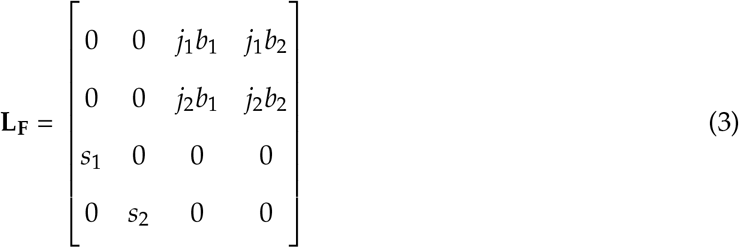

where the rows and columns of the transition matrix follow the same phenotype and age structure as the state vector 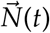 (equation (2)).

In contrast, the dynamic condition model allows individuals to transition between pheno-types throughout their life. At each time step, we assume that an individual transitions from phenotype *x* to phenotype *y* with probability *f*_*x*→*y*_. For example *f*_2→1_*s*_2_ would represent the fraction of juveniles born in phenotype 2 that survive (with probability *s*_2_) and then transition to phenotype 1 as adults (with birth rate *b*_1_ at the next timestep). In the case of only two age classes there is only one possible age transition for surviving individuals (age 1 to 2), however in the next section we allow this transition probability to also depend on an individual’s age. The transition matrix for a genotype with vital rates that can vary throughout life is therefore 

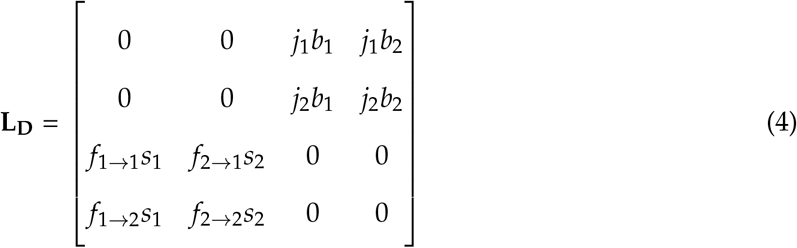

This model reduces to uncorrelated condition (fig. 1C), where phenotypes are randomly determined at each age, when *f*_*x*→*y*_ = *f*_*y*→*y*_ = *f*_*y*_ for all phenotypes *x* and *y* (i.e., the probability of transitioning into a given phenotype is the same for all individuals of a given age).

Consider the long-term growth rates of the fixed and dynamic matrices in equations (3) and (4). Both of these are given by the leading eigenvalue of the transition matrices above 

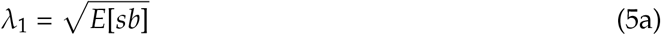

Here, *E*[*sb*] can be thought of as the expected value of the adult birth rate, accounting for all of the possible life history “paths” (phenotypic transitions and corresponding survival probabilities) that allow an individual to survive to the adult class in one of the two phenotypes. Recalling that by definition the covariance of two variables (here, *s* and *b*) is equal to the expected value of their product minus the product of their expected values, i.e., *Cov*(*s, b*) = *E*[*sb*] − *E*[*s*]*E*[*b*], equation (5a) can be rewritten as

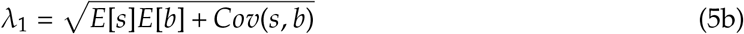

Here *E*[*s*] gives the expected survival probability to the adult age class, while *E*[*b*] gives the expected birth rate, now independent of survival but still taken over all of the possible phenotypic transitions that an individual could have experienced throughout its life course.

Importantly, equation (5b) reveals that a genotype’s fitness will be different from what would be expected with independent vital rates any time that there is a nongenetic covariance among survival to the adult class and the birth rate within that class. The only case where this covariance will be 0 is when the vital rates are stochastic throughout the lifespan, such that individuals transition randomly to any phenotype at any time (uncorrelated condition with *f*_1→1_ = *f*_2→1_ and *f*_1→2_ = *f*_2→2_). Correspondingly, the covariance will be strongest when vital rates are fixed throughout the lifespan. Thus, given the same age structure, individual vital rates, and transition probabilities, a genotype’s fitness will be greater than expected with uncorrelated condition when survival probabilities and birth rates are positively related and lower than its fitness with uncorrelated vital rates when survival probabilities and birth rates are negatively related.

The relationship between the type of individual heterogeneity, the type of covariance, and the population growth rate can be explained by the fraction of individuals that have the greatest survival probability as juveniles and the greatest birth rate as adults. These high condition individuals contribute disproportionately to a genotype’s fitness, through their own survival and through their offspring. For example, without loss of generality, assume that phenotype 1 individuals have the highest survival probability and the highest birth rate in a genotype with a positive covariance among vital rates. Comparing the two extremes at the stable phenotype distribution (defined below), the proportion of individuals in this phenotype category with fixed condition is 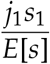 and with uncorrelated condition is 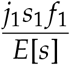. Assuming there is heterogeneity (0 < *f*_1_ < 1), the fraction of high condition individuals will always be greater with fixed condition than with uncorrelated condition, explaining why population growth rates are higher under fixed condition when vital rates positively covary. Analogous calculations show that if there is a trade-off between survival probabilities and birth rates (*Cov*(*s, b*) < 0), the proportion of high condition individuals (for example, phenotype 1 for survival but phenotype 2 for reproduction) will always be greater with uncorrelated condition than with fixed condition (Supplemental material S1).

The influence of the covariance among vital rates on population dynamics is underscored by considering an example population where juveniles keep their phenotype to the adult age class with probability *τ* (fixed condition), and correspondingly, are assigned a random phenotype with probability 1 − *τ* (uncorrelated condition). The long-term population growth rate is then a weighted average of the growth expected for fixed (*E*[*sb*]) and uncorrelated condition (*E*[*s*]*E*[*b*])

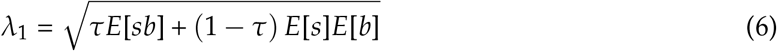

The sensitivity of this growth rate with respect to *τ* is

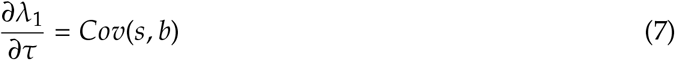

Accordingly, when the covariance among survival probabilities and birth rates is positive, an increase in the persistence of vital rates throughout an individual’s life course (*τ*) will increase the population growth rate; negative covariances have the opposite effect.

#### The Full Model

We now consider *ρ* vital rate phenotypes and *ω* ages. This general model allows reproduction from individuals of any age and it allows individuals to survive up to age *ω*. At a given time, an individual belongs to one of *ρ* · *ω* states. Thus, the state vector is 

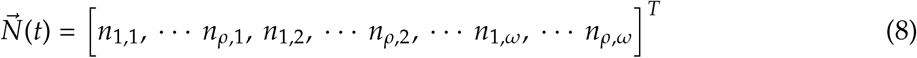

Further, let 

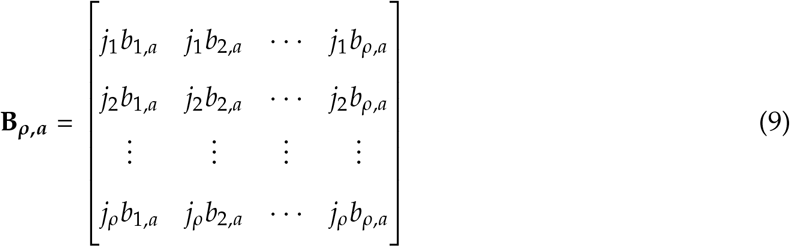

be a matrix of the potential birth rates from individuals of age *a* to any of the *ρ* phenotypes, where *j*_*x*_ represents the probability that an adult of age *a* gives birth to offspring that are in phenotype *x* at age 1. Assume that 

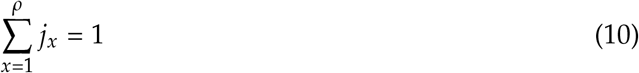

so that all offspring from individuals of age *a* must be born into one of the *ρ* phenotypes and there is no death of newborns from the transition process. These transition probabilities do not depend on the phenotype or age class of the parent, a limiting assumption that we make in order to collapse the matrices without loss of information about the dynamics as described later in this section. Note, however, that there will often be some kind of parent-offspring correlation (e.g., due to maternal effects, nongenetic inheritance, or correlated environments between parents and offspring (Bonduriansky and Day 2009; Danchin et al. 2011; Hernández et al. 2020)), which must be treated using the full phenotype-age matrix to preserve the full population dynamics and eigenstructure (although the leading eigenvalue and stable age distribution can still be obtained by collapsing the matrix as described below, see Hooley 2000 and Coste et al. 2017).

To incorporate heterogeneity throughout the lifespan, suppose that the potential survival and transition probabilities for individuals of age *a* and phenotype *x* are given by the matrix 

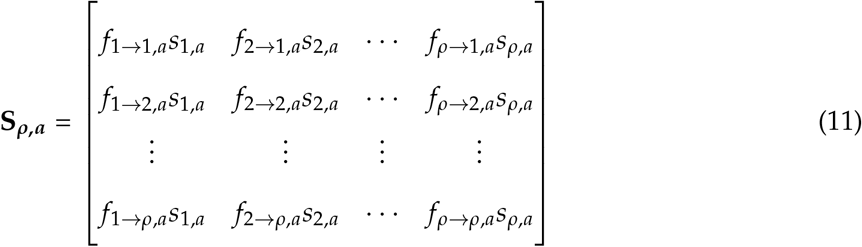

Here, *f*_*x*→*y*,*a*_ represents the probability that a surviving individual of phenotype *x* at age *a* transitions into phenotype *y* by age *a* + 1. These transition probabilities can depend on an individual’s current and future age and phenotype. For notational simplicity, we include only the subscript for the current age in these transition probabilities, since the next age is directly determined by the current age *a* (individuals cannot skip or move backwards in age). Further, assume that 

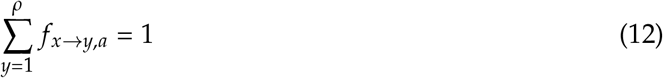

and so there is no loss of surviving individuals due to the transition process. This matrix allows transitions between phenotypic categories at arbitrary rates, including the two extreme types of individual heterogeneity (fixed and uncorrelated condition). If *f*_*x*→*y*,*a*_ = 1 when *y* = *x* and is 0 otherwise, then the phenotypes are permanent throughout life (fixed condition), while if *f*_*x*→*y*,*a*_ = *f*_*y*→*y*,*a*_ = *f*_*y*,*a*_ for all phenotypes *x* and *y* then the condition is uncorrelated, with phenotypes randomly drawn at each age. An intermediate choice allows the phenotype to be correlated from one age to the next, with probability *τ* of remaining in the same phenotype and probability (1 − *τ*) *f*_*y*,*a*_ of being assigned a phenotype at random in the next time step (here *τ* measures the strength of the phenotypic correlation between age classes).

Under these assumptions, the *ρ*-phenotype *ω*-age transition matrix takes the form 

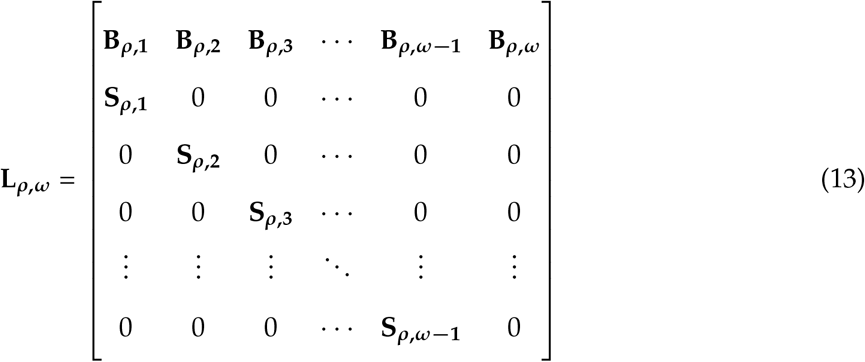

where **L**_***ρ***,***ω***_ is a square matrix of size *ρ* · *ω*-by-*ρ* · *ω* and each 0 in **L**_***ρ***,***ω***_ is a *ρ* · *ρ* block of 0’s. Readers familiar with IPMs will recognize this structure (Easterling et al. 2000; Ellner and Rees 2006) and could easily extend our model to compare the effects of different kinds of heterogeneity when phenotypic categories are continuous.

The generalized transition matrix in equation (13) can be used to predict ecological and evolutionary dynamics more accurately than matrices that oversimplify the true number of pheno-types and age classes in a population. However, the matrices will often be very large if they account for the number of phenotypes and age classes found in real populations. For example, if there are only five potential phenotypes for survival, five for birth rates, and two age classes in a population, the matrix accounting for individual heterogeneity will already be of size (5 · 5 · 2) *×* (5 · 5 · 2) = 50 *×* 50.

Here we present a novel matrix collapsing method to reduce the *ρ*-phenotype *ω*-age matrix to an *ω*-age matrix that contains a single phenotype for each age class (fig. 1D). Accordingly, these collapsed matrices take the same form as traditional Leslie matrices. This technique was inspired from previous work that shows how to collapse population projection matrices across age or stage classes (Enright et al. 1995; Hooley 2000; Salguero-Gómez and Plotkin 2010). Although these previous matrix collapsing methods typically only guarantee that the long-term growth rate and stable age distribution of the original matrix are captured by the collapsed matrix (Coste et al. 2017; Hooley 2000; with the method of Bienvenu et al. 2017 also preserving reproductive values), we prove that the full solution of the original matrix model is captured by our method for the kinds of matrices considered here, where phenotypes among newborns are chosen at random (Appendix 1; see examples in Supplemental material S2). This means that the collapsed matrix captures both the transient and asymptotic dynamics of age-structured populations with such nongenetic heterogeneity in vital rates, after a short transition period of at most *ω* time steps (see Supplemental *Mathematica* document for examples) and that all eigenvalue and eigenvector pairs of the collapsed system completely characterize the solution of the full system (see Appendices). Briefly, our application of matrix collapsing (termed “phenotypic collapsing”) merges the rows and columns corresponding to the *ρ* phenotypes within each age class (see Appendix 1 for the full recipe). The phenotypes are weighted by the proportion of individuals in age *a* and phenotype *x* that are expected from offspring born *a* time steps ago, considering all phenotypic transitions and survival rates that could have occurred in between (*ψ*_*x*,*a*_, see Appendix 1; Appendix 2). Note that these weights give the stable phenotypic distribution for each age class after at most after *ω* time steps. A major advantage of phenotypic collapsing is that it does not require that the system is at a stable age distribution, which is different from classic matrix collapsing that uses the stable age distribution to determine the collapsing weights (Enright et al. 1995; Hooley 2000). Additionally, phenotypic collapsing does not rely on calculating and weighting by the eigenvectors, requiring only the total probability of all possible paths from birth to survival at age *a* with phenotype *x* (a total that is described by a simple matrix power, see Appendix 1). A full worked example of collapsing a transition matrix with two phenotypes and two or three age classes to a transition matrix with a single-phenotype for each age class is found in Supplemental material S2.

Define *l*_*a*_ as the probability of surviving up to age *a*. The collapsed *ω*-age matrix can then be shown to equal 

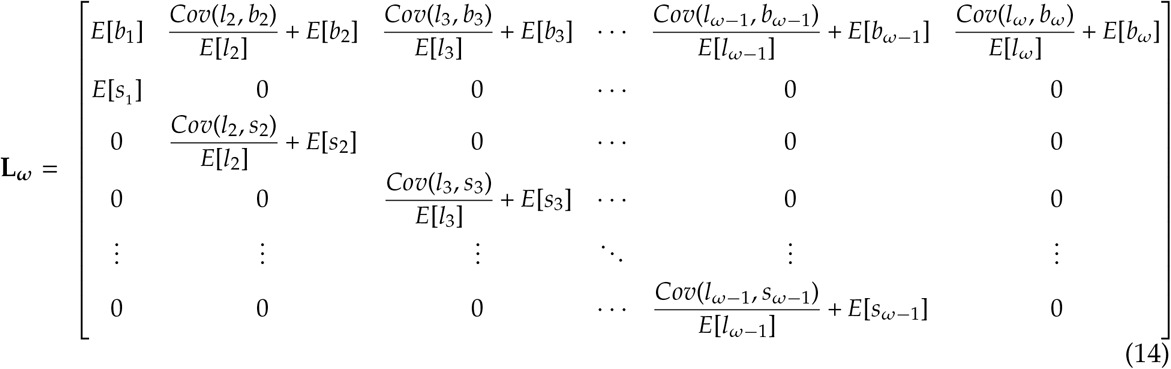

where the matrix entries are calculated using the phenotypic collapsing recipe provided in Appendix 1. The covariances (*Cov*) are taken between all of the possible phenotypic histories that an individual could have experienced to survive up to a given age *a* (*l*_*a*_), and the birth rate (top row) or survival probability (subdiagonal) at that age. The expectations of the vital rates at age *a, E*[*s*_*a*_] and *E*[*b*_*a*_], are taken over all of the possible phenotypic transitions that an individual could have experienced up to age *a*, independent of survival along the way (i.e., with *ψ*_*x*,*a*_ = *f*_*x*,*a*_ for each phenotype *x* at age *a* > 1 and *j*_*x*_ for the newborns, see Appendices for proof and Supplemental material S2 for examples). These are the expected values of the vital rates that would be obtained if the current phenotype did not depend on any previous phenotype (uncorrelated condition), and so can be calculated as an average taken only over the phenotypic distribution at that age. The expectation *E*[*l*_*a*_] is the expected probability of surviving to age *a*, considering all of the possible phenotypes and corresponding survival probabilities that an individual could have experienced throughout its life course from age 1 to *a* (i.e., with *ψ*_*x*,*a*_ depending on previous transition and survival probabilities, see Appendices and examples in Supplemental material S2).

The collapsed matrix **L**_***ω***_ reveals that when the current survival and birth rates are independent of the life history path that an individual took to get there (i.e., the heterogeneity is uncorrelated and phenotypes are randomly determined at each age), all covariances are 0 and the effective vital rates can be calculated as a weighted average considering only the vital rates and transition probabilities assigned to each age (ignoring any previous transition or survival probabilities, such that *ψ*_*x*,*a*_ = *f*_*x*,*a*_ or *j*_*x*_ as mentioned above). All else being equal, increasing the covariance between survivorship to age *a* and the survival probability or birth rate at age *a* will increase the effective vital rates in equation (14). Consequently, when there is a positive covariance among vital rates over the lifespan, the long-term growth rate will be higher than predicted using only the average vital rates at each age, with the opposite holding when there are trade-offs among vital rates.

Another important observation is that the collapsed vital rates in equation (14) are simply the average vital rates of individuals surviving to age *a* (Appendix 2). Considering the collapsed matrix in this way underscores its value for providing straightforward predictions of ecological and evolutionary dynamics even when there are many heterogeneity categories in a population, however it overlooks the influence of the covariance among vital rates on these dynamics (see Discussion below).

The collapsed matrix can be used to analyze populations with vital rate heterogeneity, with-out requiring extra matrix dimensions to account for this individual variation. Indeed, when applied to matrices in the form of equation (14), we find that phenotypic collapsing preserves the complete solution of the original matrix for all times *t* ≥ *ω*, such that the eigenvalue and eigenvector pairs of the collapsed system completely characterize the full system (Appendix 1; see examples in Supplemental material S2). We have also found that all non-zero eigenvalues and their associated eigenvectors from the original *ρ*-phenotype *ω*-age matrix are captured by the collapsed matrix, as long as the population has reached a stable phenotype distribution (this is proven for cases with non-repeating eigenvalues in Supplementary material S2 and will be proved more generally in a forthcoming article). This property holds because the weights that are used to collapse do not depend on the chosen eigenvalue or eigenvector once the distribution of phenotypes is constant from one generation to the next. In our case, this condition is satisfied because there is no phenotypic memory from parents to offspring, and so the stable phenotype distribution is achieved within a single full lifetime as the distribution of phenotypes at birth is fixed and independent of the population vector. Thus, while matrix collapsing using the right eigenvector (here, the stable age *and* phenotype distribution) was already known to preserve the asymptotic population growth rate (*λ*_1_) and its associated right eigenvector (stable age distribution) (Hooley 2000) and in some cases can also maintain the associated left eigenvector (reproductive values) (Coste et al. 2017) of the original full size matrix, here we show that phenotypic collapsing also preserves the full solution and so the transients as well for certain kinds of matrices (e.g., when the heterogeneity is “reset” at some age, as in equation (13)).

### Numerical Analysis - Multiple Genotypes

#### Methods

To explore the ecological and evolutionary consequences of nongenetic heterogeneity in vital rates when multiple clonal genotypes are co-evolving, we compliment our single-genotype analyses with randomly drawn genotypes and calculate the mean asymptotic growth rate and evolvability (rate of evolution by natural selection) among them. By definition, the mean asymptotic growth rate is the mean fitness for a genotype, however in many empirical studies mean fitness is approximated by the average number of surviving offspring per parent. For clarity, we use the term “asymptotic growth rate” here. Note that these analyses assume that each genotype has reached its stable age and phenotypic distribution when the mean growth rate and evolvability are calculated.

Intuitively, the rate of evolution in the asymptotic population growth rate can be calculated as 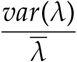 where the mean and variance are taken across the growth rates of each genotype and the resulting rate is in units of population growth (*λ*). However, this quantity measures the response to selection (“evolvability”) on an absolute scale, and so it cannot be used to compare rates of evolution across populations and environments with different growth rates. For example, a population where individuals reach reproductive maturity after a single time step (“early recruitment,” below) would be expected to grow faster than a population where maturity takes ten time steps (“late recruitment,” below). Here we instead use *relative* evolvability as the appropriate measure of the rate of evolution. As shown by Houle (1992), relative evolvability is calculated as the response to selection, standardized by the mean asymptotic growth rate (i.e., 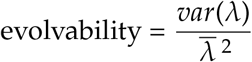). This approach provides a dimensionless measure of the variance in geno-type growth rates and thus allows comparison across populations with different life histories and with different strengths and types of nongenetic (co)variance. Note that relative evolvability is influenced both by the mean and variance in growth rates; hence we provide results for the unstandardized variance in Supplemental material (fig. S1; fig. S2). Further, note that evolvability predicts only short term changes in the asymptotic population growth rate relative to its current value; in the long run, the genotype with the largest growth rate is expected to go to fixation in this density-independent model. In addition to influencing evolvability, we also find that when cohort selection is strong enough (e.g., when there is late recruitment, below) the strength of nongenetic heterogeneity influences which genotype has the highest growth rate (i.e., the rank order of the genotypes depends on the heterogeneity in phenotypes each exhibits).

For simplicity, this numerical analysis considers only the two extreme types of nongenetic individual heterogeneity: fixed and uncorrelated condition. We also use a small number of age classes and vital rate phenotypes to facilitate comparison with the single-genotype analysis, however the model is easily extended to incorporate more complex population structures.

Recalling that the sign of the covariance among individual vital rates can influence genotype fitness, we consider positive, zero, and negative genetic and nongenetic covariances among survival probabilities and birth rates. The general modeling procedure is described below and full details can be found in the Supplemental material S3. All calculations were performed in the R statistical environment (ver. 4.3.1; R Core Team 2023). The full R code is provided in Forsythe et al. (2024) https://datadryad.org/stash/share/8Yh2oLMu4BXHhrurxgKfd1DEZjM2BqvteGyeyCQ4tz8.

A genotype is defined by its set of mean vital rates. To ensure constant and biologically realistic genotype means (i.e., all survival probabilities between 0 and 1 and all birth rates greater than or equal to 0), we randomly drew vital rates from a bivariate uniform distribution. The genetic covariance was incorporated with a positive, negative, or 0 covariance in the variance-covariance matrix.

To keep the numerical analyses at least somewhat tractable and compatible with our analytical results, we assume that each genotype contains only two possible phenotypes for each vital rate. The strength of the nongenetic variance determines how far the individual survival probabilities and birth rates fall above (phenotype 1) or below (phenotype 2) their genotypic mean. We allow the strength of the nongenetic variance to range from 0 to 5 times as much as the genetic variance, which is held constant throughout all calculations. Note that because the phenotypic means are held constant (at the genotypic mean), the only variable that changes across our numerical analyses is the strength of the nongenetic heterogeneity.

The final step to parameterize the single-genotype matrices is to calculate the transition probabilities for each phenotype (*f* and *j* above). For simplicity we assume that the transition probabilites are independent of age and either completely determined by previous vital rates (fixed condition) or independent of any previous vital rates (uncorrelated condition), however this assumption could be relaxed, allowing partial persistence of the phenotype from age-to-age as in equation (11). The transition probabilities are calculated as the fraction in each quadrat under a bivariate normal distribution that is centered at the genotypic mean vital rates and with non-genetic covariance given by one of the three structures defined earlier: zero, positive, or negative.

Given the individual vital rates and the transition probabilities, we can parameterize the fixed and uncorrelated condition matrices for a population with any number of non-reproductive and reproductive ages. Here, we consider mean fitness and evolvability for a population with one juvenile age and one adult age (early recruitment into the breeding population) and for a population with ten juvenile ages and one adult age (late recruitment). The early recruitment example matches the simple example in the theoretical analysis, but with a total of four phenotypes to account for all possible combinations of two survival probabilities and two birth rates. To facili-tate comparison, we use the same vital rates, transition probabilities, and number of juvenile and adult ages to parameterize the fixed and uncorrelated condition matrices.

#### Early recruitment: Higher nongenetic covariance increases mean fitness, but slows rate of evolution

First consider the results with early recruitment (fig. 2). Recall from equation (14) that the fitness of a genotype with uncorrelated condition depends only on the expected values of the agespecific survival probabilities and birth rates at the age they are experienced, while the fitness of a genotype with fixed condition depends on these expected values *and* the nongenetic covariance among these vital rates and all of the possible ways that an individual could have survived to experience them. All else being equal, positive nongenetic covariances increase the average vital rates in the transition matrix with fixed condition, and this is expected to translate into an increase in the population growth rate, while negative covariances have the opposite effect. These results scale up to determine the mean asymptotic growth rate among genotypes, which for uncorrelated condition is constant regardless of the nongenetic covariance because the expected values of the vital rates are held constant in all numerical analyses (fig. 2A). The results for fixed condition are also exactly predicted from the theoretical analyses, where a stronger positive covariance among vital rates results in a higher mean asymptotic growth rate across genotypes, while a stronger negative covariance reduces the mean asymptotic growth rate across genotypes (fig. 2A). When there is no nongenetic covariance, the fixed and uncorrelated condition matrices are the same and the mean asymptotic growth rate depends only on the expected values of the vital rates at each age. Importantly, the effects of nongenetic heterogeneity on the mean asymptotic growth rate can be larger than the effects of genetic heterogeneity, even when the variation in both is the same.

**Figure 2.**
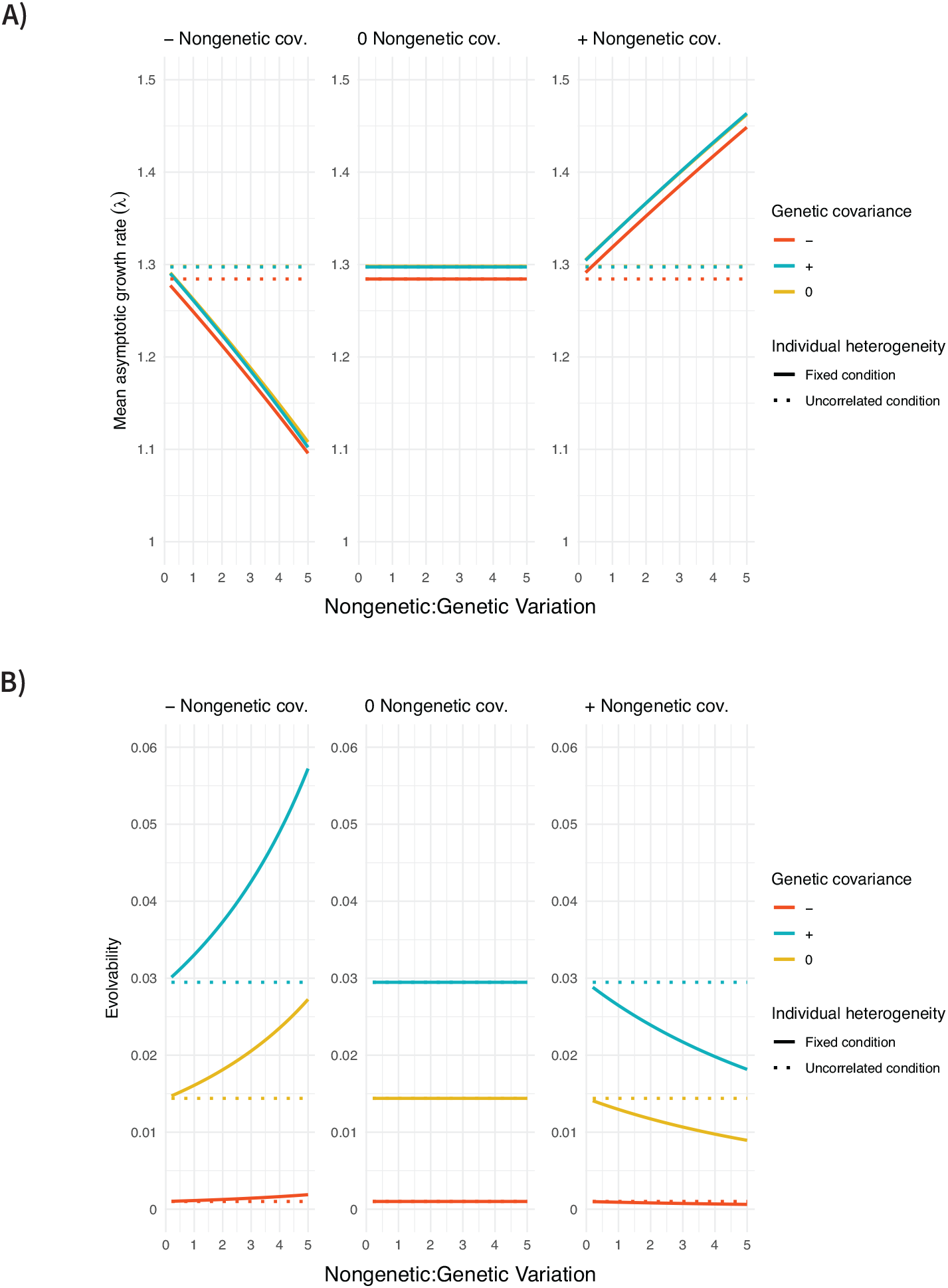
Impact of genetic and nongenetic heterogeneity on **A)** mean asymptotic growth rate (given by the leading eigenvalue, *λ*_1_) and **B)** evolvability (given by the standardized variance in asymptotic growth rates) across 1000 randomly drawn genotypes when there is early recruitment to the breeding population. The mean asymptotic growth rate and evolvability (y-axes) are functions of the relative strength of nongenetic variation (x-axes), the sign of the genetic and nongenetic covariance (line colours and panels, respectively), and the type of individual heterogeneity (line types). Note that the average fitness across all genoytpes (i.e., the average number of surviving offspring per parent) is the same in all cases, but the asymptotic growth rate given by the leading eigenvalue of the corresponding transition matrices rises with positive genetic covariance (blue lines) and, for fixed condition, with positive nongenetic covariance (right panel).

The sign of the genetic covariance has a consistent effect on evolvability, measured as the variance in asymptotic growth rates standardized by their mean squared (fig. 2B). Positive genetic covariances result in the highest evolvability because genotypes with a high mean survival probability and birth rate will replicate much faster than genotypes with a low mean survival probability and birth rate. Likewise, the lowest evolvability occcurs with negative genetic covariances because genotypes with high mean values of one vital rate will generally have low mean values of the other. Different genotypes will therefore tend to be more similar in their asymptotic growth rates, compared to positive covariances that allow for clearly superior and inferior genotypes. In further agreement, evolvability from the calculations with no genetic covariance falls in between the results from positive and negative genetic covariance.

Turning to nongenetic heterogeneity, when vital rates are uncorrelated the relative strength of nongenetic variation has no influence on evolvability (fig. 2B). Although the variation in survival and birth rates increases with the relative amount of nongenetic heterogeneity, it does so equally up (high fitness phenotypes) and down (low fitness phenotypes), and so the variance in asymptotic growth among genotypes should not change because the asymptotic growth rate depends only on the expectations of vital rates (which remain unchanged). This result also holds for fixed condition with no nongenetic covariance among vital rates. However, when the non-genetic covariance is positive, increasing the relative strength of nongenetic heterogeneity always reduces evolvability with fixed condition. Indeed, the “lucky” individuals with high survival probabilities and high birth rates dominate more strongly as the strength of nongenetic heterogeneity increases; this results in part due to cohort selection removing low condition phenotypes (Kendall et al. 2011; Vaupel et al. 1979) and in part due to an increase in vital rates for all high condition phenotypes. Accordingly, the lucky phenotypes for the different genotypes are more similar to one another in fitness and evolvability declines. The opposite effect occurs for negative nongenetic covariances among vital rates, where increasing the strength of nongenetic variation always increases the evolvability with fixed condition. With a negative nongenetic covariance, phenotypes tend to have either a high survival probability and low birth rate, or a low survival probability and high birth rate, causing the phenotypic heterogeneity to cancel out to some extent (i.e., all individuals are mediocre) and accentuating genotypic differences in asymptotic growth rates. Thus, unintuitively, the population evolves more quickly but towards a lower mean asymptotic growth rate as the strength of nongenetic heterogeneity is increased with a negative nongenetic covariance.

#### Late recruitment: Fixed condition increases mean fitness, but slows rate of evolution

We next consider longer life spans, with reproduction only after 10 time steps (late recruitment). All else being equal, late recruitment reduces the asymptotic growth rate compared to early recruitment because individuals must survive longer before contributing to the population via reproduction (fig. 3A). Again, if the nongenetic heterogeneity is uncorrelated (dashed lines), the mean asymptotic growth rate is determined directly by the expectations of the survival probabilities and birth rates at each age, and so there is no influence of the sign of the nongenetic covariance. By contrast, the asymptotic growth rates are substantially higher when vital rates are fixed across the life span, increasingly so as the nongenetic heterogeneity in vital rates rises. In this case, there is a stronger frailty effect and the majority of phenotypes that survive long enough to reproduce are the “lucky” phenotypes with a high survival probability. With a positive nongenetic covariance, this phenotype is also the one that tends to have high birth rates, allowing for the strongest increase in asymptotic growth rate with the strength of nongenetic heterogeneity. With negative nongenetic covariance, however, the individuals that are in the high survival probability phenotype will have the lowest birth rates, and so there is a slightly weaker increase in genotype fitness. The impact of the nongenetic covariance between survival and birth rates is, however, much more modest with late recruitment, because survival rates across all age classes are the same (i.e., survival at age *a* and *a* + 1 are completely positively correlated).

**Figure 3.**
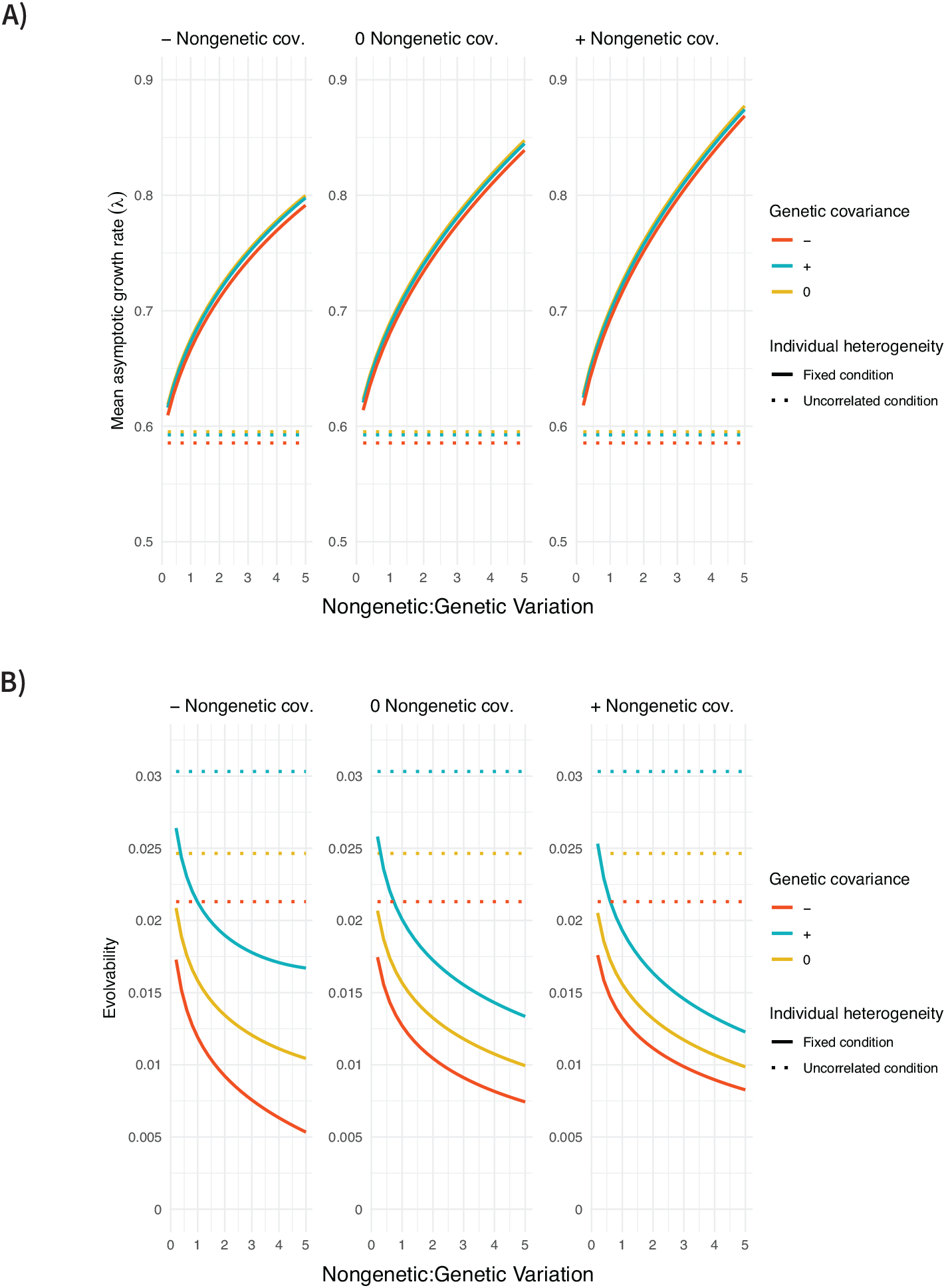
Impact of genetic and nongenetic heterogeneity on **A)** mean asymptotic growth rate (given by the leading eigenvalue, *λ*_1_) and **B)** evolvability (given by the standardized variance in asymptotic growth rates) across 1000 randomly drawn genotypes when there is late recruitment to the breeding population. The mean asymptotic growth rate and evolvability (y-axes) are functions of the relative strength of nongenetic variation (x-axes), the sign of the genetic and nongenetic covariance (line colours and panels, respectively), and the type of individual heterogeneity (line types). Note that the average fitness across all genoytpes (i.e., the average number of surviving offspring per parent) is the same in all cases, but the mean asymptotic growth rate given by the leading eigenvalue of the corresponding transition matrices rises with the relative strength of nongenetic heterogeneity, unless the vital rates are reset randomly each time step (uncorrelated condition, dashed lines).

As with early recruitment, the sign of the genetic covariance has a consistent effect on evolvability, where all else being equal, positive genetic covariances result in the highest evolvability and negative genetic covariances in the lowest evolvability (fig. 3B). Likewise, increasing the strength of nongenetic heterogeneity has no influence on the rate of evolution by natural selection when vital rates are uncorrelated over the lifespan. However, when the vital rates are fixed at birth, evolvability decreases as the relative strength of nongenetic variation increases; this is another consequence of only the few “lucky” individuals with high survival probabilities surviving long enough to reproduce, with genotypic differences being relatively modest.

This relatively simple example with uniformly distributed vital rates emphasizes the effects of nongenetic individual heterogeneity because all parameters are held constant across the numerical analyses, except the amount of nongenetic variation. However, vital rates will very rarely follow such a perfect distribution in natural populations. In more realistic situations, the processes generating nongenetic heterogeneity are likely to affect both the mean and variance of each vital rate, which will complicate the results depending on that relationship (e.g., Plard et al. 2016).

## Discussion

Here we introduced phenotypic collapsing as a method that allows phenotypic heterogeneity in survival probabilities and birth rates to be captured within a traditional Leslie matrix, while preserving the full solution of the original full-scale model. Previous studies of matrix collapsing use a technique that typically enforces the same leading eigenvalue and stable age distribution as the original matrix (Hooley 2000; Salguero-Gómez and Plotkin 2010), however it need not capture the left eigenvector associated with the leading eigenvalue (reproductive values) (Bienvenu et al. 2017; Coste et al. 2017) nor the remaining eigenstructure. Furthermore, by instead collapsing over the stable phenotype distribution as we do here (i.e., using the fraction of newborns that reach age *a* in phenotype *x* as collapsing weights, rather than the stable age distribution), we provide additional insight into the role that covariances between vital rates play in ecological and evolutionary dynamics. Our model is general in the sense that it makes no assumptions about the factors that generate and maintain variation in these vital rates, it allows for any number of age classes and phenotype categories, and it can be applied to a wide range of species, however it does require that the phenotype distribution is reset at birth, with randomly drawn juvenile phenotypes regardless of the parent’s age or phenotype. While our numerical analyses focus on the cases where vital rates are set at birth (fixed condition) or can vary randomly at anytime (uncorrelated condition), the theoretical analyses provide further insight by allowing for arbitrary phenotypic transitions across the lifespan, as will often be the case in natural populations (Orzack et al. 2011; Plard et al. 2012, 2018).

Previous works support our result that nongenetic variation in vital rates can affect ecological and evolutionary dynamics. For example, Cressler et al. (2017) used an individual-based model to show that regardless of the sign of the nongenetic covariance among vital rates, rates of evolution by natural selection (calculated as the variance in genotypic growth rates) will always be slower with higher relative strengths of nongenetic individual heterogeneity. This model assumed fixed condition and that individuals had to survive at least ten time steps to reproduce, and so their findings correspond to our special case of fixed condition and late recruitment (fig. 3B). Our study provides more general insight, however, as we find that evolvability can instead sometimes increase with the strength of nongenetic heterogeneity (as in the case where there is a trade-off among vital rates and early recruitment (fig. 2B)) or remain unchanged (e.g., with uncorrelated condition).

Importantly, we find that the sign of the nongenetic covariance among individual survival probabilities and birth rates has a predictable influence on the long-term growth rate of a geno-type. A similar effect was recognized by Stover et al. (2012), who incorporated fixed condition in birth and death rates into a logistic model with density-dependence in reproduction. The authors observed that a positive correlation among these vital rates (assumed to be independent of age) always increases the low-density population growth rate and equilibrium population size, while a negative correlation decreases both measures. Likewise, Kendall et al. (2011) found that long-term population growth rates increase with the magnitude of heterogeneity in survival rates, which were held constant throughout life, creating a positive *Cov*(*l*_*a*_, *s*_*a*_) in equation (14) (heterogeneity in birth rates had no effect in their model without a parent-offspring correlation, however, because it was uncorrelated with survival rates, so *Cov*(*l*_*a*_, *b*_*a*_) were zero by assumption for all ages *a*). Here we expanded on the results of Kendall et al. (2011) and Stover et al. (2012) by explicitly considering genetic and nongenetic components of individual heterogeneity in vital rates in an age-structured model with an arbitrary number of phenotype categories and by exploring their consequences on evolvability. We also compare results to a model of dynamic condition, where individuals can change phenotypes throughout life.

The main result from our theoretical analysis showed that the long-term growth rate of a genotype depends on how vital rates do or do not vary throughout the life course. At one extreme, if vital rates vary randomly throughout the life course (uncorrelated condition), we find that population dynamics depend only on the average vital rates within an age class and are independent of any previous phenotypic history allowing suvival to that age (i.e., all covariances in equation (14) are 0). This finding corresponds to a study from Fox and Kendall (2002), which determined that if there is variability in vital rates but they are independent and identically distributed, then the influence of this heterogeneity on population dynamics disappears and the outcome matches what would be expected if all individuals had the same, average vital rate. All else being equal, we find that positive nongenetic covariances among survival up to an age class and the survival or birth rate at that age translate to greater genotypic fitness than expected if vital rates varied randomly throughout the life course, while trade-offs have the opposite effect (equation (14)). Under a positive covariance, individuals that have high survival probabilities also have high birth rates, and these “lucky” individuals substantially increase genotype fitness throughout their lives. In natural populations of the parlour palm and rainforest trees, for example, high condition individuals contribute up to 2.4 times as much to population growth rates as low condition individuals (Jansen et al. 2012; Zuidema et al. 2009). However, trade-offs are also widespread in nature and they occur when individuals that have a high survival probability pay a cost with a low birth rate or vice versa (e.g., Dahlgren et al. 2016; Gould et al. 2018; Lemaître et al. 2015). Trade-offs are expected to provide higher fitness when life histories are dynamic because individuals that have a high survival probability at one time can potentially move among phenotypes and have a high birth rate at the next (Supplemental material S1).

Whether the covariances are positive or negative, the model predicts that vital rate hetero-geneity will have a stronger effect in the life-history stages where phenotypes are longer-lived (leading to stronger covariances). For example, if transitions are dynamic early in life, but fixed (highly correlated) later (perhaps after development), then the covariance among vital rates would be expected to have a much larger effect at older ages. This result corresponds with a recent study from Bliard et al. (2024), which found that populations with longer lifespans are more sensitive to changes in the strength of vital rate trade-offs because cohort selection has more opportunities to take place than in populations with shorter lifespans. Our finding also implies that even when covariances among survival probabilities and birth rates are expected to be strong, their impacts on population dynamics may be masked by frequent transitions among phenotypes. Similarly, Lee et al. (2017) found that reproductive autocorrelations (as would be produced in our models of fixed or dynamic condition) have a stronger effect on the demographic variance and stochastic population growth rate when females have a low probability of switching reproductive status as they age. Our framework could therefore explain why previous studies have found that heterogeneity has only a weak effect on the variance among individual longevity and lifetime reproductive success compared to stochasticity in vital rates (Hartemink and Caswell 2018; Jenouvrier et al. 2018; Snyder and Ellner 2018; Steiner and Tuljapurkar 2012): the effects of heterogeneity are hidden by frequent transitions occurring under stochasticity.

These effects of the sign of the nongenetic covariance among vital rates on a single genotype scale up to determine the mean asymptotic growth rate among multiple co-evolving genotypes (fig. 2A). In the simple case of two age classes (“early recruitment”), the effect of the covariance among vital rates on population dynamics is most distinct and the mean asymptotic growth rate is greater for fixed condition when nongenetic covariances are positive, but greater for uncorrelated condition when nongenetic covariances are negative. In agreement, simulations from an individual based model of fixed condition (Cressler et al. 2017) showed that increasing the strength of nongenetic variation increases genotype growth rates when nongenetic covariances are positive, while negative covariances have the opposite effect. Our analytical results thus explain how the sign and magnitude of the nongenetic covariance among vital rates determines genotype growth rates in Cressler et al. (2017).

In contrast, if individuals have longer development times (“late recruitment”), we observed that fixed condition always results in higher mean asymptotic growth rates than uncorrelated condition and that this difference increases with the strength of nongenetic heterogeneity (fig. 3A). Fixed condition means that individuals that are born with a high survival probability are guaranteed to maintain that high survival probability throughout their life. If instead vital rates are dynamic, individuals that have a high survival probability at one time may transition and have a low survival probability at the next; it is therefore less likely that individuals will survive to reproduce and genotype fitness will generally be lower than with fixed condition. This result emphasizes the importance of heterogeneity for long-lived species, where maturity generally occurs later in life and so cohort selection within a generation has more time to remove the weaker individuals and leave only those with a sufficiently high survival probability to reproduce (Bliard et al. 2024; Kendall et al. 2011; Lloyd et al. 2020; Vaupel and Yashin 1985). The nuances of the type of individual heterogeneity, the vital rate distributions, the sign of the (non)genetic covariance, and the number of age classes can generate a wide range of ecological and evolutionary dynamics across populations.

Interestingly, we found only a small influence of the sign of the genetic covariance on the mean asymptotic growth rate among genotypes. This result is also similar to the results of Cressler et al. (2017), which found that the mean genotype growth rate is sensitive to the sign of the nongenetic covariance among vital rates, but largely insensitive to the genetic covariance structure. Overall, these findings emphasize that long-term growth rates and evolvability can depend equally, if not more, strongly on nongenetic (co)variances as genetic.

Our numerical analyses reveal that populations may face a trade-off between growing faster (i.e., having the greatest mean asymptotic growth rate) and evolving faster (fig. 2; fig. 3). Indeed, if life-histories are fixed condition, we found that increasing the magnitude of nongenetic heterogeneity benefits the mean asymptotic growth rate when it reduces evolvability and vice-versa. Thus, populations consisting of individuals with permanent vital rates may have an advantage for growth in a constant environment but may not be able to adapt as quickly in the face of environmental change. Future work should ask whether such environmental fluctuations influence the type of individual heterogeneity that is found across real populations.

Historically, a substantial barrier to incorporating individual heterogeneity in theoretical models has been the cost of added complexity when individuals cannot be considered identical. Here, we applied phenotypic collapsing to reduce population projection matrices with an arbitrary number of phenotypes and ages to a standard Leslie matrix, while maintaining the complete solution of the original full-scale matrix, after a short delay (of at most *ω* time steps) so that no individuals remain alive from the initial population (whose configuration was arbitrary). This work builds on previous theory that has developed collapsing methods that capture all asymptotic properties of the matrix (e.g., Bienvenu et al. 2017; Coste et al. 2017), but unlike our approach, do not necessarily preserve the transient dynamics for all times *t* ≥ *ω*. Another advantage of phenotypic collapsing is that the collapsing weights are calculated using only matrix powers and the distribution of juvenile phenotypes (see Appendix 1), which are generally much easier to compute for large matrices than the eigenvectors required in previous methods. This collapsed matrix can be used in future theoretical and empirical studies to consider variation in mortality and birth rates without sacrificing model accuracy.

The collapsed matrix also provides insight about when among-individual variation in vital rates can be ignored in studies of population dynamics to avoid unnecessary complexity (see Appendix 2 below). Equation (A2_1) shows that empirical researchers can accurately predict population dynamics using only the average age-specific vital rates, even when there is non-genetic heterogeneity in those vital rates within each age class. As long as researchers ensure random sampling and as long as the phenotypic distribution at each age is stable (e.g., the projection matrix is of the form of equation (13) and has been stable since birth), all of the ways to survive up to an age class and then have a particular survival probability or birth rate at that age will be accounted for in the expected values of the vital rates. In support, previous simulations that used an IPM to account for variation in survival and fecundity found no influence of this heterogeneity on the asymptotic population growth rate when compared to the same population analyzed with an IPM ignoring such variation (Vindenes and Langangen 2015). This method will be sufficient for many population-level studies, however it will fail if the goal of the analysis is to measure variability among life histories and its influence on demographic parameters. The matrix in equation (14) directly incorporates all of the life history paths that an individual could follow throughout its life course and highlights the importance of the covariance between an individual’s history and its current phenotype in determining population dynamics.

Here we have shown that nongenetic individual heterogeneity in vital rates can influence genotype fitness as well as the mean long-term growth rate and evolvability in populations of isolated genotypes. We restricted our analyses to density-independent age-structured models to isolate the consequences of nongenetic (co)variation as much as possible, however we expect that nongenetic heterogeneity will generally affect ecological and evolutionary dynamics. One limitation of our method is that we assumed no parent-offspring correlations in phenotype for a given genotype. Relaxing this assumption would allow the model to account for various degrees of nongenetic heritability and for potential negative correlations in the vital rates of parents and offspring (e.g., due to density effects or spatial heterogeneity), although it is important to note that the collapsed population projection matrices would be different from those that we have shown here and would not necessarily maintain the characteristic polynomial, reproductive values, or transient population dynamics. To add further biological realism, influences of environmental stochasticity (Engen et al. 2007; Sæther and Engen 2015), and continuous variation in vital rates (Plard et al. 2018; Vindenes and Langangen 2015) could also be considered. While the precise effects of the type of individual heterogeneity and nongenetic (co)variance would be less clear, these models may provide more accurate predictions of growth rates and evolvability in real populations. Our results emphasize the importance of not only considering among-individual variation in vital rates, but also the structure of that variation (e.g., genetic or nongenetic, fixed or dynamic condition, positive or negative covariance), when making predictions about ecological and evolutionary processes.

## Acknowledgments

We would like to thank Dr. Ophélie Ronce for feedback that improved this version of the manuscript and Dr. Christophe Coste for insights about matrix collapsing. This work was supported by the Natural Sciences and Engineering Council of Canada (graduate scholarships to ABF (CGS-M and CGS-D) and Discovery Grants to TD (NSERC RGPIN-2022-03362) and SPO (NSERC RGPIN-2022-03726)) and the University of British Columbia (Four Year Doctoral Fellowship to ABF).

## Statement of Authorship

ABF, TD and WAN conceptualized the work and developed the models. ABF performed model analysis, developed the R code, and carried out the numerical analyses with substantial input from SPO and TD. SPO wrote the first draft of the *Mathematica* code (edited and finalized by ABF). ABF wrote the original draft of the manuscript with input from SPO and TD and all authors contributed to editing and revisions.

## Appendix 1: A proof for phenotypic collapsing

*Notation and set up*: The model is 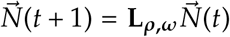 where 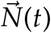 is a vector whose elements *n*_*x*,*a*_ indicate the number of individuals in phenotype *x* at age *a* and 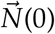 is the initial condition at time *t* = 0. We can decompose **L**_***ρ***,***ω***_ as **L**_***ρ***,***ω***_ = **F** + **V** where **F** is a square ferility matrix of dimension *ρ* · *ω* and contains the first *ρ* rows of **L**_***ρ***,***ω***_ and zeros eleswhere, and **V** = **L**_***ρ***,***ω***_ − **F**. Notice that **V** is a block subdiagonal matrix that tracks the survival of individuals. **V** consists of *ω* − 1 blocks and so it is nilpotent of degree *ω* (i.e., **V**^*τ*^ = 0 for all integral powers *τ* ≥ *ω*). We focus on cases where the probability that a juvenile starts in phenotype *x* is *j*_*x*_, independent of the parental age, phenotype, or genotype. The probability distribution of these newborns is given by the vector

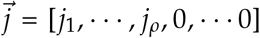 whose first *ρ* elements give the fraction of offspring born in phenotype *x* and whose remaining *ρ* · (*ω* − 1) elements are zeros. Define 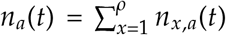 to be the total number of individuals of age *a* at time *t*, 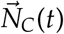 with elements *n*_*a*_(*t*) to be the *ω*-dimensional vector describing the total number of individuals in each age class, and 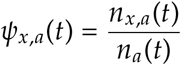 to be the fraction of age *a* individuals at time *t* that have phenotype *x*.

*Theorem*: Consider the *ρ* · *ω* dimensional initial value problem 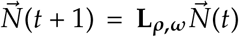 with initial vector at time *t* = 0 given by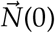. The solution 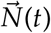 for time *t* ≥ *ω* is completely described by the solution of the reduced, *ω* dimensional initial value problem 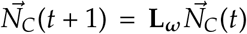 with initial vector at time *t* = *ω* given by 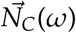 and where the elements of **L**_***ω***_ are block averages over time-invariant values of *ψ*_*x*,*a*_ (*t*) defined by 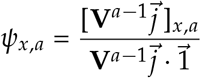 where 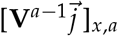 is the *x, a*th entry of the vector given by 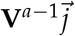 and 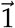 is a *ρ* · *ω* dimensional vector of ones.

*Proof* : Consider the collapsed matrix **L**_***ω***_ obtained by averaging the elements of **L**_***ρ***,***ω***_ over the phenotypic distribution *ψ*_*x*,*a*_(*t*) for each age class *a*. Only if *ψ*_*x*,*a*_(*t*) is time invariant will 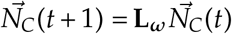 represent a linear system of equations involving the *ω* entries of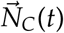, describing the total number of individuals in each age class. Here we will show that, for times *t* ≥ *ω* these *ψ*_*x*,*a*_(*t*) are in fact time invariant and given by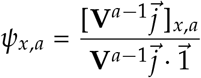. Thus for *t* ≥ *ω*, the solution of the original model lies in the *ω*-dimensional subspace spanned by the eigenvalues and eigenvectors of the collapsed model. In other words, for any time *t* ≥ *ω*, the matrix **L**_***ω***_ obtained from phenotypic collapsing completely predicts the dynamics of the original full matrix, even if the stable age distribution has not yet been reached.

We begin by writing the solution 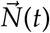 in renewal form as 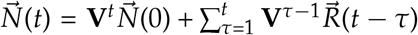 for *t* ≥ 1 where 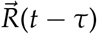 is a vector whose first *ρ* elements are the total recruitment of offspring of phenotype *x*, by all individuals present at time *t* − *τ*, and whose remaining elements are zeros. Because **V** is nilpotent of degree *ω*, for times *t* ≥ *ω* we have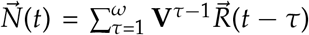. Furthermore, because there is no memory of parent phenotypes,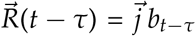 where *b*_*t*−*τ*_ is a scalar giving the total number of new recruits of age one individuals of all phenotypes by all individuals present at time *t* − *τ*. Therefore, 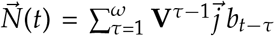 for *t* ≥ *ω*. Notice that the time dependence, *t*, appears only in *b*_*t*−*τ*_. Finally, it can be checked that because of the form of **V** and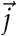, the terms 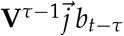 in the summation of 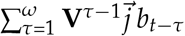 are each a vector with zeros everywhere except for the block corresponding to age *τ*. Therefore, 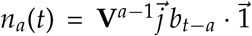 and 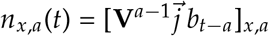. Thus after simplifying, 

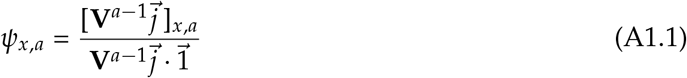

which is independent of time *t*.

*Recipe*: Define **A** to be an *ω ×* (*ρ* · *ω*) matrix with rows for each age class *a* (in order from 1 to *ω*) that contain *ρ* ones in the slot for age class *a* and zeros elsewhere. Define **B** to be a (*ρ* · *ω*) *× ω* matrix with columns for each age class *a* (in order from 1 to *ω*) containing the collapsing weights, *ψ*_*x*,*a*_ for each age *a* and zeros elsewhere. The collapsed matrix is then given by 

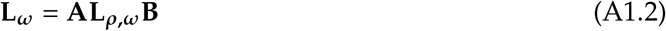

Notice that the matrices **A** and **B** follow the same structure as **P** and **Q** (respectively) in the collapsing method of Hooley (2000), but with entries that collapse over phenotypes instead of age classes and with collapsing weights *ψ*_*x*,*a*_ determined by the stable phenotype distribution rather than the stable age and phenotype distribution. A major advantage of phenotypic collapsing is that these weights (*ψ*_*x*,*a*_, equation (A1.1)) can be calculated using only matrix powers (of the survival matrix, **V**) and the distribution of juvenile phenotypes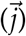. Previous collapsing methods on the other hand (e.g., Bienvenu et al. 2017; Coste et al. 2017; Hooley 2000) require an eigenanalysis of the orginial, full-scale matrix to calculate the collapsing weights (typically determined by the dominant eigenvalue and eigenvector pair), and these computations become challenging to compute analytically or even numerically for very large matrices.

## Appendix 2: Some properties of phenotypic collapsing

First, notice that for all times *t* ≥ *ω* the collapsing weights given by *ψ*_*x*,*a*_ are simply the probability that an individual is in phenotype *x* at age *a*, given that it survived to age *a*. Thus we can also write 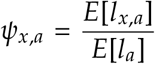 where *E*[*l*_*x*,*a*_] is the expected probability of lifetime survival, from a random juvenile phenotype leading up to an age *a* individual in phenotype *x*, calculated over the possible juvenile distribution. Specifically, let 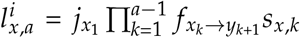 represent one of several possible paths *i* that an individual born with phenotype *x*_1_ could take to survive and express phenotype *x*_*a*_ = *x* at age *a*, considering all of the phenotypes *x*_*k*_ that the individual experienced between birth and age *a*. The sum over all of these paths 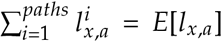 gives a weighted average of all potential phenotypic transitions leading an individual to phenotype *x* at age *a* (where the weights are determined by the transition probabilities that determine how likely each path is); this summation is an alternative mathematical expression of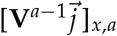. The sum over all paths and phenotypes 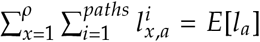 gives a weighted average of all of the ways that an individual can survive to age *a*, considering all of the phenotypes *x* that it could express at that age, which is equivalent to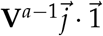. These entries *ψ*_*x*,*a*_ of the stable phenotype distribution can be put together into the vector 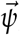, which gives the stationary proportion of individuals in each phenotype *x* for each age *a*. As it turns out, the stationary phenotypic proportions for each age given in 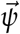 are the same as would be obtained at the stable age *and* phenotype distribution (given by the right eigenvector associated with the leading eigenvalue of **L**_***ρ***,***ω***_), however our methods do not require that the stable age distribution has been reached.

Our application of matrix collapsing uses this stable phenotype distribution to collapse all of the phenotypes within each age class to a single, effective phenotype for that age class. The weightings for each phenotype *x* at age *a* are given by *ψ*_*x*,*a*_ as defined above and in Appendix 1. The effective vital rates are then given by 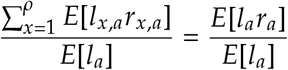 where *r* is one of the two vital rates (survival probability or birth rate), and *E*[*l*_*a*_*r*_*a*_] gives an expected value of the vital rate at age *a*, considering all of the potential phenotypes that could be expressed at age *a* and all of the possible life histories (i.e., phenotypic transitions and survival probabilities) that an individual could have experienced as it survives up to that age.

Recalling that for any variables *x* and *y* the expected value *E*[*xy*] can be written in terms of the covariance, *E*[*xy*] = *E*[*x*]*E*[*y*] + *Cov*(*x, y*), we now seek to define what these expectations should be if there were no covariance. If the current vital rates did not depend on previous vital rates, we could obtain the phenotypic distribution at age *a* as above, but ignoring previous survival

rates. Hence, we define 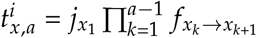 to be the probability of the path *i* for an individual born in phenotype *x*_1_, considering all of the phenotypes *x*_*k*_ that it transitions through to express phenotype *x*_*a*_ = *x* at age *a*, as if differences in survival rates at previous ages did not affect the current vital rates. Setting the survival probabilities to one in the survival matrix **V** (as defined in Appendix 1), we get a matrix that we call **V**_**T**_ and 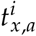 can be calculated from the entries of 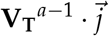 (note that the transition probability of all possible paths from birth to age *a* sum to one for each age *a*, i.e.,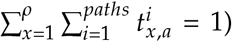. Defining 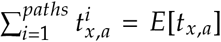 and 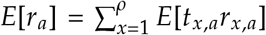 the collapsed *ω*-age matrix can then be written as in equation (14).

As an alternative, we note that equation (14) can also be written as 

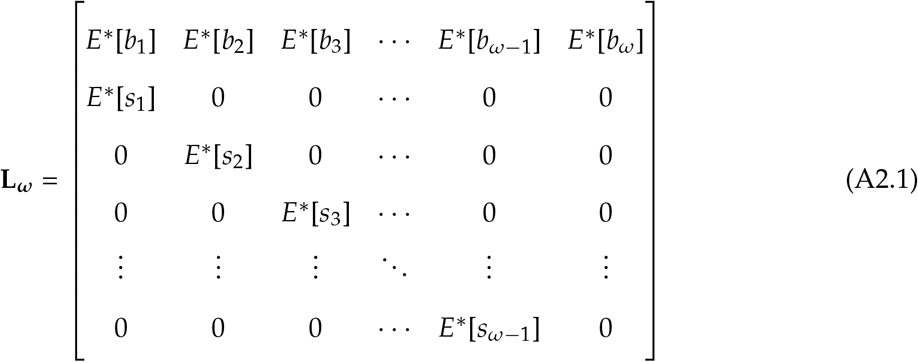

where the expectations, *E*^*\**^, are taken over over all of the possible phenotypic histories that an individual could have experienced to survive to age *a*, accounting for both the transition and survival probabilities encountered along the way, that is, defining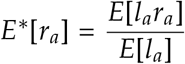. This form of the collapsed matrix shows that empirical estimates of age-specific vital rates are accurate, even when there is heterogeneity in the vital rates within each age class, as long as there is random sampling and the population is at the stable phenotype distribution, as expected if the vital rates have remained constant for *t* ≥ *ω* timesteps. This alternative form of the collapsed matrix, however, hides the impact of phenotypic heterogeneity and the covariance among vital rates on the growth of the population, as revealed by equation (14).

As argued here and in Appendix 1 (see also worked examples in Supplemental material S2) the stable phenotypic distribution 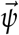 is attained in at most a single full lifetime, once no individuals remain alive from the initial population (*t* ≥ *ω*). Consequently, as long as the juvenile distribution does not depend on the parental age and phenotype, the collapsed matrix, alongside the time invariant phenotypic distribution 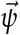, fully describes the future state of the system for all *t* ≥ *ω*. The solution of the original model with the matrix **L**_***ρ***,***ω***_ is therefore simply a linear combination of the eigenvalues and eigenvectors associated with the collapsed matrix **L**_***ω***_ (see Supplemental material S2 for an example). In a forthcoming article we will derive the explicit relationship between the eigensolutions of the full and the reduced system and explore more generally the kinds of matrix models that can be simplified using phenotypic collapsing. This implies that the characteristic polynomials of the collapsed and original *ρ*-phenotype *ω*-age matrix are proportional to one another and exactly equal when put in Euler-Lotka form (see Supplemental material S2 for an example). The Euler-Lotka equation (Euler 1760; Sharpe and Lotka 1911) is given by setting the characteristic polynomial equal to zero and rearranging terms such that 

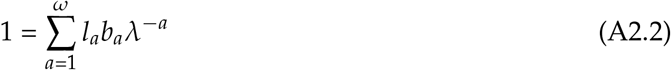

where *l*_*a*_ is the proportion of individuals surviving to age *a* and *b*_*a*_ is the average number of offspring born to an individual of age *a*, assuming that the population is at its stable age distribution. This equation reveals that the average lifetime reproductive output for individuals is the sum of all age-specific contributions to fitness, via births and survival.

The characteristic polynomial for the collapsed matrix with heterogeneity is 

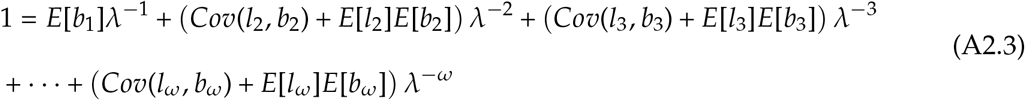

This equation further emphasizes the influence of the covariance among vital rates on population growth when individual life histories have at least some phenotypic “memory.” All else being equal, the average contribution of an age class to lifetime reproductive output can be higher (positive covariance), lower (negative covariance), or the same (no covariance) as if the phenotypes were randomly assigned.

While the phenotypic distribution described by 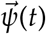 holds immediately for all *t* ≥ *ω*, the stable age distribution will often take several generations to be approached. A major advantage of phenotypic collapsing is that it does not assume that the population is at a stable age distribution and therefore the collapsed matrix **L**_***ω***_ can still describe transient dynamics as the age structure stabilizes (Appendix 1; see examples in Supplemental material S2). Here we briefly consider the stable age distribution with phenotypic collapsing (see worked examples in Supplemental material S2). The entries of the stable age distribution for age *a* in the collapsed matrix **L**_***ω***_ are given by the expected value of survival to age *a, E*[*l*_*a*_], as described by summing all of the paths from birth to age *a* accounting for all possible phenotypic transitions (equivalently by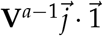).

Notice that these entries of the stable age distribution of the collapsed matrix are simply the sum of the entries of the stable age and phenotype distribution from the original matrix **L**_***ρ***,***ω***_

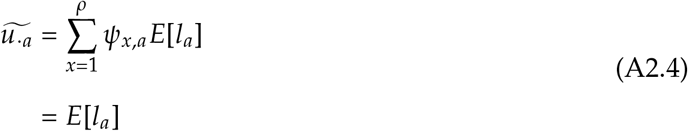

Equation (A2_4) falls in agreement with previous studies of matrix collapsing, which find that the entries of the stable age or stage distribution of the original matrix (in this case, the entries of the stable age and phenotype distribution of **L**_***ρ***,***ω***_) can be summed over the collapsed classes to calculate the entries of the stable (st)age distribution for the collapsed matrix (Hooley 2000; Salguero-Gómez and Plotkin 2010).

Previous studies of matrix collapsing (which use the stable (st)age distribution as collapsing weights) guarantee that the dominant eigenvalue and its associated right eigenvector from the original matrix will be captured in the collapsed matrix. However, in contrast to phenotypic collapsing, even the left eigenvector associated with the dominant eigenvalue (which gives the reproductive value of each class) is only captured in particular cases with this method. For example, a study from Coste et al. (2017) found that matrix collapsing over the stable state distribution can capture reproductive values for matrices where the “contributions from all states (or future states) are equally broken down with regards to the soon-to-be-grouped states” (Coste et al. 2017). The matrices that we use here fit this condition because the heterogeneity is “reset” at the newborn stage when there is no memory of parental phenotypes (Coste pers. comm.); this implies that the age-specific reproductive values are the same before and after collapsing, if they are weighted by the stable phenotype distribution. The age-specific reproductive values, 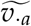, of the collapsed *ω*-age matrix are therefore given by 

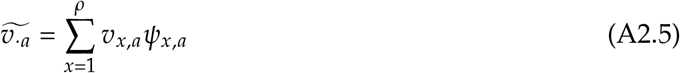

where the reproductive values of the original *ρ*-phenotype *ω*-age matrix, *v*_*x*,*a*_, indicate the asymptotic contributions of an individual in each age-and-phenotype specific class to future generations, relative to individuals in other classes (see worked examples in Supplemental material 2).

The reproductive values of each age class in the collapsed matrix are therefore proportional to the reproductive values of the *ρ* phenotypes that were merged to form that age class from the original *ρ*-phenotype *ω*-age matrix. Weighting by *ψ*_*x*,*a*_ ensures that the contribution of each age class to future generations is scaled by the proportion of individuals that are expected in each phenotype category in the long run. Equation (A2_5) is also in agreement with the form of reproductive values from Bienvenu et al. (2017), who developed a different collapsing technique (“genealogical collapsing”) that always captures the asymptotic population growth rate, as well as both its associated right and left eigenvectors from the original full matrix. Note, however, that the weightings applied to the reproductive values by Bienvenu et al. (2017) assume that the population is at its stable stage distribution (given by the right eigenvector associated with the dominant eigenvalue), whereas the weightings we use only require that the population is at a stable phenotype distribution, which is obtained once *t* ≥ *ω* and all individuals in the population develop from juveniles drawn from the same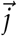. Furthermore, with phenotypic collapsing, the general solution and therefore the full nonzero eigensystem is captured, not just the leading eigenvalue and its associated eigenvectors. Thus, the future dynamics of the system are fully predicted, without assuming that the system is at the stable age distribution.

### Code Availability

All R code for performing numerical analyses and generating figures is available in the Dryad Digital Repository (https://datadryad.org/stash/share/8Yh2oLMu4BXHhrurxgKfd1DEZjM2BqvteGyeyCQ4tz8). Forsythe et al. 2024).

## Footnotes

1 This type of heterogeneity is conceptually similar to what previous studies have referred to as “individual stochasticity” (e.g., Forsythe et al. 2021; van Daalen and Caswell 2020), “luck” (e.g., Snyder and Ellner 2018, 2022), or “dynamic heterogeneity” (e.g., Steiner et al. 2010; Tuljapurkar et al. 2009), however an important distinction is that here we use “uncorrelated condition” to refer to random variation among individual vital rates, not individual fates.

