## Supplemental Material for "Variety is the spice of life: nongenetic variation in life histories influences population growth and evolvability"

June 1 2024

1. Department of Zoology and Biodiversity Research Centre, University of British Columbia, Vancouver, BC V6T 1Z4, Canada;

2. Department of Biology, Queen's University, Kingston, Ontario K7L 3N6, Canada;

3. Department of Mathematics and Statistics, Queen's University, Kingston, Ontario K7L 3N6, Canada.

### S1. An explanation for why nongenetic covariances determine genotype fitness.

Consider the two-phenotype two-age example in the main text with the two extreme types of heterogeneity: fixed condition and uncorrelated condition. We would like to know the proportion of individuals that have the highest survival probability as juveniles *and* the highest birth rate as adults, since these are the "lucky" individuals that will contribute the most to population growth throughout their lives. Note that these fractions could be calculated from $\psi_{x,a}$ (see Appendices), which gives the probability that an individual is in phenotype $x$ at age $a$, given that it survived to age $a$ and that we have reached the stable phenotype distribution. For a population with a maximum of two ages, the stable phenotype distribution is achieved after just two timesteps given any starting population vector (Appendix 1). Here we instead arrive at the same conclusion by tracking a cohort of individuals throughout their lives, still assuming that the stable phenotype distribution has been reached.

The stable age and phenotype distribution, given by the right eigenvector associated with the dominant eigenvalue (our measure of the asymptotic population growth rate) and normalized to sum to one, is

$$\vec{u}_F = \frac{1}{\sqrt{E[sb]} + E[s]} \left[ j_1 \sqrt{E[sb]},\ j_2 \sqrt{E[sb]},\ j_1 s_1,\ j_2 s_2 \right] \tag{S1}$$

for fixed condition and

$$\vec{u}_D = \frac{1}{\sqrt{E[s]} + \sqrt{E[b]}} \left[ j_1 \sqrt{E[b]},\ j_2 \sqrt{E[b]},\ f_1 \sqrt{E[s]},\ f_2 \sqrt{E[s]} \right] \tag{S2}$$

for uncorrelated condition. Note that these vectors give the same proportion of individuals in each phenotype as we would expect if only the stable phenotype distribution had been reached (Appendix 1). Thus, at the stable phenotype distribution, the proportion of phenotype 1 juveniles is $\frac{j_1 \sqrt{E[sb]}}{j_1 \sqrt{E[sb]} + j_2 \sqrt{E[sb]}} = j_1$ for fixed condition and $\frac{j_1 \sqrt{E[b]}}{j_1 \sqrt{E[b]} + j_2 \sqrt{E[b]}} = j_1$ for uncorrelated condition. Likewise, the proportion of phenotype 2 juveniles is $j_2$, regardless of the type of individual heterogeneity.

An individual's phenotype is permanent throughout life under fixed condition, and so the fraction of the phenotype 1 juveniles that survive to become phenotype 1 adults is determined by the survival probability for phenotype 1. Given that we started with $j_1$ juveniles in phenotype 1, the proportion surviving to become adults is $j_1 s_1$. Note that these adults will also have the birth rate given by $b_1$. Similarly, the surviving individuals with the vital rate set $(s_2, b_2)$ is calculated as $j_2 s_2$, and there are no individuals that transition among phenotypes. Scaling by the total fraction of surviving individuals of either phenotype, the proportion of phenotype $i$ individuals is $\frac{j_i s_i}{E[s]}$ where $i = 1$ or 2. With uncorrelated condition however, juveniles can change phenotypes as they survive to become adults. The proportion of the total surviving individuals with vital rates $(s_1, b_1)$ is then $\frac{j_1 s_1 f_1}{E[s]}$ and the fraction of individuals with vital rates $(s_2, b_2)$ is $\frac{j_2 s_2 f_2}{E[s]}$. Note that the total fraction of surviving individuals is the same as with fixed condition ($E[s]$), even though the fractions of "lucky" individuals are different. Since individuals can transition between phenotypes with uncorrelated condition, we also have a nonzero proportion of individuals with vital rates $(s_1, b_2)$ and $(s_2, b_1)$. These proportions are $\frac{j_1 s_1 f_2}{E[s]}$ and $\frac{j_2 s_2 f_1}{E[s]}$, respectively.

Without loss of generality, suppose that $s_1 > s_2$ and $b_1 > b_2$, such that there is a positive covariance among vital rates and phenotype 1 individuals are the most fit. The fraction of individuals with a high survival probability *and* high birth rate is higher for fixed condition than for uncorrelated condition, since as long as there is heterogeneity $0 < f_x < 1$ and therefore $j_1 s_1 > j_1 s_1 f_1$. Hence, a positive covariance among vital rates means that there will be a higher fraction of these "lucky" individuals if the individual heterogeneity is fixed condition than if it is uncorrelated. These individuals increase population growth rates considerably throughout their lives, explaining why positive covariances result in faster population growth rates with fixed condition than uncorrelated condition.

On the other hand, if $s_1 > s_2$ but $b_1 < b_2$, there is a negative covariance (trade-off) among vital rates. If the individual heterogeneity is fixed condition, there are no individuals with high survival probabilities and high birth rates; individuals are either stuck in phenotype 1 with a high survival probability and low birth rate, or in phenotype 2 with a low survival probability and high birth rate. However, since individuals may transition to a new phenotype with uncorrelated condition, the proportion of individuals with a high survival probability and high birth rate is $\frac{j_1 s_1 f_2}{E[s]}$. Again as long as there is heterogeneity, $0 < f_x < 1$, and now the fraction of "lucky" individuals is higher if the individual heterogeneity is uncorrelated condition than if it is fixed condition, explaining why population growth rates are fastest with uncorrelated condition when there is a trade-off among vital rates.

### S2. Matrix collapsing procedure.

Phenotypic collapsing uses the stable phenotype distribution as weights to collapse vital rate phenotypes within age classes of a transition matrix to a single, effective phenotype for each age class (see Appendices in the main text). This technique is inspired by an existing collapsing method, “Ergodic Flow Preserving” (EFP) merging (Hooley 2000; Enright et al. 1995; Coste et al. 2017), however importantly phenotypic collapsing maintains the complete solution of the original matrix for any time $t \geq \omega$ (see Appendices and examples in the attached *Mathematica* notebook), whereas EFP merging is only guaranteed to capture the dominant eigenvalue and its associated right eigenvector (the asympotic population growth rate and stable age distribution, respectively). Thus with phenotypic collapsing all eigenvalues and eigenvectors of the reduced system completely characterize the full system for any time $t \geq \omega$. In a forthcoming article we will further derive the explicit relationship between the eigensolutions of the full and the reduced system (see attached *Mathematica* notebook for a proof for cases with non-repeating eigenvalues).

Here we provide a detailed example of phenotypic collapsing and in the attached *Mathematica* notebook we show that it preserves the characteristic polynomial (in Euler-Lotka form), and it captures all non-zero eigenvalues and their associated right and left eigenvectors from the original full scale matrix for the case of two phenotypes with two and three age classes. For this example, consider the transition matrix for a two-age model with fixed condition, as given by equation (3) in the main text. We will use the recipe outlined in Appendix 1 to collapse this matrix to a single-phenotype two-age matrix, while preserving the all non-zero eigenvalues and their associated right and left eigenvectors of the original two-phenotype two-age matrix. The characteristic polynomial (simplified to Euler-Lotka form) of the original matrix is

$$1 = E[sb]\lambda^{-2} \tag{S3}$$

where

$$E[sb] = j_1 s_1 b_1 + j_2 s_2 b_2 \tag{S4}$$

Solving for $\lambda$ in (S3) gives the eigenvalues of $\mathbf{L_F}$, and the largest of these eigenvalues is the population growth rate

$$\lambda_F = \sqrt{E[sb]} \tag{S5}$$

The stable age and phenotype distribution of the matrix $\mathbf{L_F}$ (standardized such that the entries sum to 1) is

$$\vec{u}_F = \frac{1}{\lambda_F + E[s]} \left[ j_1\lambda_F,\ j_2\lambda_F,\ j_1 s_1,\ j_2 s_2 \right] \tag{S6}$$

where $E[s] = j_1 s_1 + j_2 s_2$. Note that $\vec{u}_F$ is given by the right eigenvector associated with the population growth rate, or can be found by iterating the full phenotype-by-age survival matrix to find the fraction of individuals surviving to age $a$ and with phenotype $x$ at that age (see Appendices and *Mathematica* notebook).

The phenotype- and age-specific reproductive values of $\mathbf{L_F}$ are given by the left eigenvector associated with the population growth rate

$$\vec{v}_F = \frac{E[s] + \lambda_F}{2\lambda_F^3} \left[ s_1 b_1,\ s_2 b_2,\ b_1\lambda_F,\ b_2\lambda_F \right] \tag{S7}$$

where $\vec{v}_F$ is standardized such that $\vec{v}_F.\vec{u}_F = 1$.

The goal here is to collapse over the two phenotypes in $\mathbf{L_F}$, so that we are left with a typical Leslie matrix with a single set of effective vital rates for each age class. To merge the phenotypes in age class 1, we need to merge rows 1 and 2 and columns 1 and 2. Likewise, to merge the phenotypes in age class 2, we need to merge rows 3 and 4 and columns 3 and 4. The first step is to define weights that represent the long run proportions of the merged age class that are expected from each of the the original phenotypes in that age class. Let $\psi_{x,a}$ be the weight for phenotype $x$ in age class $a$. Then from Appendix 1

$$\psi_{x,a} = \frac{[\mathbf{V}^{a-1}\vec{j}\,]_{x,a}}{\mathbf{V}^{a-1}\vec{j} \cdot \vec{1}} \tag{S8}$$

where $\mathbf{V} = \begin{bmatrix} 0 & 0 & 0 & 0 \\ 0 & 0 & 0 & 0 \\ s_1 & 0 & 0 & 0 \\ 0 & s_2 & 0 & 0 \end{bmatrix}$ is the matrix capturing survival of individuals and $\vec{j} = \begin{bmatrix} j_1 & j_2 & 0 & 0 \end{bmatrix}$ is a vector giving the probability distribution for the phenotypes of the newborn offspring and with zeros elsewhere. Note this is equivalent to $\psi_{x,a} = \dfrac{E[l_{x,a}]}{E[l_a]}$ for all $t \geq \omega$ (where here $\omega = 2$ is the maximum age here) as shown in Appendix 2. Furthermore, the stable age and phenotype distribution could be used to compute the same probability, $\psi_{x,a} = \dfrac{u_{x,a}}{\sum_{x=1}^{\rho} u_{x,a}}$ where $u_{x,a}$ is the proportion of individuals in phenotype $x$ and age class $a$ at the stable phenotype distribution (given by the $x, a$th entry of $\vec{u}_F$), and where there are $\rho$ phenotypes in each age class. Using any of these approaches gives

$$\begin{aligned} \psi_{1,1} &= j_1 \\ \psi_{2,1} &= j_2 \\ \psi_{1,2} &= \frac{j_1 s_1}{E[s]} \\ \psi_{2,2} &= \frac{j_2 s_2}{E[s]} \end{aligned} \tag{S9}$$

as the weights for this two-phenotype two-age example, where as above $E[s] = j_1 s_1 + j_2 s_2$. Using the recipe in Appendix 1, we can then define the matrices $\mathbf{A}$ and $\mathbf{B}$

$$\mathbf{A} = \begin{bmatrix} 1 & 1 & 0 & 0 \\ 0 & 0 & 1 & 1 \end{bmatrix} \tag{S10}$$

$$\mathbf{B} = \begin{bmatrix} \psi_{1,1} & 0 \\ \psi_{2,1} & 0 \\ 0 & \psi_{1,2} \\ 0 & \psi_{2,2} \end{bmatrix} \tag{S11}$$

The collapsed matrix $\mathbf{L}_\omega$ is then given by $\mathbf{L}_\omega = \mathbf{A}\mathbf{L}_{\rho,\omega}\mathbf{B}$, which for this example simplifies to

$$\mathbf{L}_\omega = \begin{bmatrix} 0 & \dfrac{E[sb]}{E[s]} \\ E[s] & 0 \end{bmatrix} \tag{S12}$$

where $\dfrac{E[sb]}{E[s]} = \dfrac{Cov(s,b)}{E[s]} + E[b]$

The characteristic polynomial of (S12) is

$$1 = E[sb]\lambda^{-2} \tag{S13}$$

which exactly matches the characteristic polynomial from (S3), as expected. Furthermore, the stable age distribution of the collapsed matrix (S12), standardized so that the entries sum to 1, is

$$\widetilde{u} = \frac{1}{\lambda_F + E[s]} \begin{bmatrix} \lambda_F, & E[s] \end{bmatrix} \tag{S14}$$

Notice that the stable age distribution of $\mathbf{L_F}$ is also maintained, in the sense that the age-specific entries of (S14) are the sum of the merged phenotype- and age-specific entries from (S6). Taking $\widetilde{u_{\cdot a}}$ to be the entries of the stable age distribution for the collapsed matrix, then mathematically

$$\widetilde{u_{\cdot a}} = \sum_{x=1}^{\rho} u_{x,a} \tag{S15}$$

as expected (Hooley 2000). The entries of the stable age distribution of the collapsed matrix can also be written in terms of the collapsing weights $\psi_{x,a}$ using the analyses above (see also Appendix 2)

$$\begin{aligned} \widetilde{u_{\cdot a}} &= \sum_{x=1}^{\rho} \psi_{x,a} \sum_{x=1}^{\rho} u_{x,a} \\ &= E[l_a] \end{aligned} \tag{S16}$$

since $\sum_{x=1}^{\rho} \psi_{x,a} = 1$ and $\sum_{x=1}^{\rho} u_{x,a} = E[l_a]$. The reproductive values of (S12) are given by

$$\widetilde{v} = \frac{\lambda_F + E[s]}{2} \begin{bmatrix} \dfrac{1}{\lambda_F}, & \dfrac{1}{E[s]} \end{bmatrix} \tag{S17}$$

where $\widetilde{v}$ is standardized such that $\widetilde{v}.\widetilde{u} = 1$. Thus, the reproductive values of $\mathbf{L_F}$ are also preserved, in the sense that

$$\widetilde{v_{\cdot a}} = \frac{\sum_{x=1}^{\rho} v_{x,a} u_{x,a}}{\sum_{x=1}^{\rho} u_{x,a}} \tag{S18}$$

where $\widetilde{v_{\cdot a}}$ are the age-specific reproductive values of the collapsed matrix, and $v_{x,a}$ are the phenotype- and age-specific reproductive values of the original matrix (Bienvenu et al. 2017). Again this equation can be rewritten in terms of the collapsing weights to match Appendix 2

$$\widetilde{v_{\cdot a}} = \sum_{x=1}^{\rho} v_{x,a}\psi_{x,a} \tag{S19}$$

We have shown that phenotypic collapsing works for an arbitrary number of phenotypes ($\rho$) and ages ($\omega$) and it works with any type of individual heterogeneity (provided that the juvenile distribution does not depend on the parental age and phenotype, see Appendix 1). The collapsed matrix is formed by merging all $\rho$ phenotypes to give a single, effective phenotype for each age class. With $\omega$ ages, the merging steps (summing rows and columns corresponding to the phenotypes in that age class) will be repeated $\omega$ times, once for each age class.

### S3. Procedures for numerical analyses.

This supplement describes the numerical analysis procedures in further detail than in the main text. All calculations were performed in the R statistical environment (ver. 4.3.1; R Core Team 2023). The full R code and output data are found at `https://datadryad.org/stash/share/8Yh2oLMu4BXHhrurxgKfd1DEZjM2BqvteGyeyCQ4tz8`. (Forsythe et al. 2024).

A genotype is defined by its set of mean vital rates ($\overline{s_g}$, $\overline{b_g}$), where $\overline{s_g}$ is the mean survival probability and $\overline{b_g}$ is the mean birth rate at each age for genotype $g$. To ensure constant and biologically realistic genotype means ($0 \leq \overline{s_g} \leq 1$ for all survival probabilities and $\overline{b_g} \geq 0$ for all birth rates), we drew vital rates from a bivariate uniform distribution. To do so, we first drew the expected vital rates for a genotype from a standard bivariate normal distribution with mean vector $(0, 0)$, a variance of one for each vital rate, and either no covariance or strong positive (0.95) or negative (-0.95) covariance. We then converted to a uniform distribution by mapping the position of the random draw along the cumulative density function (CDF) for each vital rate. These uniform draws for the genotypic mean survival probabilities and birth rates were then scaled so that $\overline{s_g}$ ranged from 0.35 to 0.65 and $\overline{b_g}$ ranged from 2.4 to 4.4. These values were selected so that individual vital rates were still biologically plausible after phenotypic heterogeneity was added (below).

To determine the phenotypes within a genotype, we first define $n = 2$ bins of values that an individual could express for each vital rate at a given time. We assume that these bins are evenly distributed within three standard deviations above and below the genotypic mean vital rate and that the midpoint, $m_i$, of a bin is representative of all potential values within its range. The individual survival probabilities, $s_i$, and birth rates, $b_i$, are calculated as

$$s_i = m_i * \sigma_s + \overline{s_g} \tag{S20}$$

$$b_i = m_i * \sigma_b + \overline{b_g} \tag{S21}$$

where $\sigma_s$ and $\sigma_b$ are the nongenetic standard deviations of the survival probability and birth rate, calculated as the square root of their variances. The nongenetic variance ($\sigma_s^2$ or $\sigma_b^2$) is given by the genetic variance from the uniform distribution of genotype means, but multiplied by a scalar ranging from 0 to 5 to explore the effects of shifting the relative amount of nongenetic and genetic variation in a population. Mathematically,

$$\sigma_v = \sqrt{h * \sigma_g^2} \tag{S22}$$

where $\sigma_v$ is the nongenetic standard deviation for the vital rate $v$, $\sigma_g^2$ is the genetic variance for that vital rate, and $h$ is the strength of nongenetic heterogeneity (a scalar ranging from 0 to 5). Note that for a given phenotype $i$, both $m_i$ and the genotype mean vital rates ($\overline{s_g}$ or $\overline{b_g}$) are constant, and so the only variable that changes is the strength of nongenetic heterogeneity. We consider all possible combinations of the $n = 2$ values of each vital rate, giving $n^2$ phenotypes of individuals within every genotype.

The final step to parameterize the single-genotype matrices is to calculate the transition probabilities, $f_x$, for each phenotype $x$. We assume these probabilities are equal for offspring and surviving individuals to keep the numerical analyses tractable. Consider a grid of all $n^2$ phenotypes of individuals in a genotype. Now, overlay that grid with a standard bivariate normal distribution, centered at the genotypic mean vital rates and with nongenetic covariance given by one of the three structures defined earlier: zero (0), positive (0.95), or negative (-0.95). The transition probabilities are calculated as the fraction under this bivariate normal distribution lying in each quadrat (both $s$ and $b$ positive, both negative, $s$ positive but $b$ negative, and $b$ positive but $s$ negative). With a positive covariance, for example, the transition probabilities will be highest for the both positive and both negative quadrats, such that "lucky" individuals who happen to have higher survival rates also tend to have higher birth rates.

Given the individual vital rates, $s_i$ and $b_i$, and the transition probabilities, $f_x$, we can parameterize the fixed and uncorrelated condition matrices (as given in the main text) for a population with any number of juvenile and adult ages. The mean asymptotic growth rate and evolvability are then numerically calculated based on the leading eigenvalues (asymptotic growth rates) from each genotype. Here we explore numerical analyses with a minimal number of phenotypes and ages and a rather restrictive transition probability ($f_x$ is the same for all ages and for uncorrelated condition is independent of the phenotype that individuals have come from) to compare outputs with our analytical results, however future work could relax these assumptions to explore the effects of heterogeneity in a broader range of population structures.

### Supplementary figures.

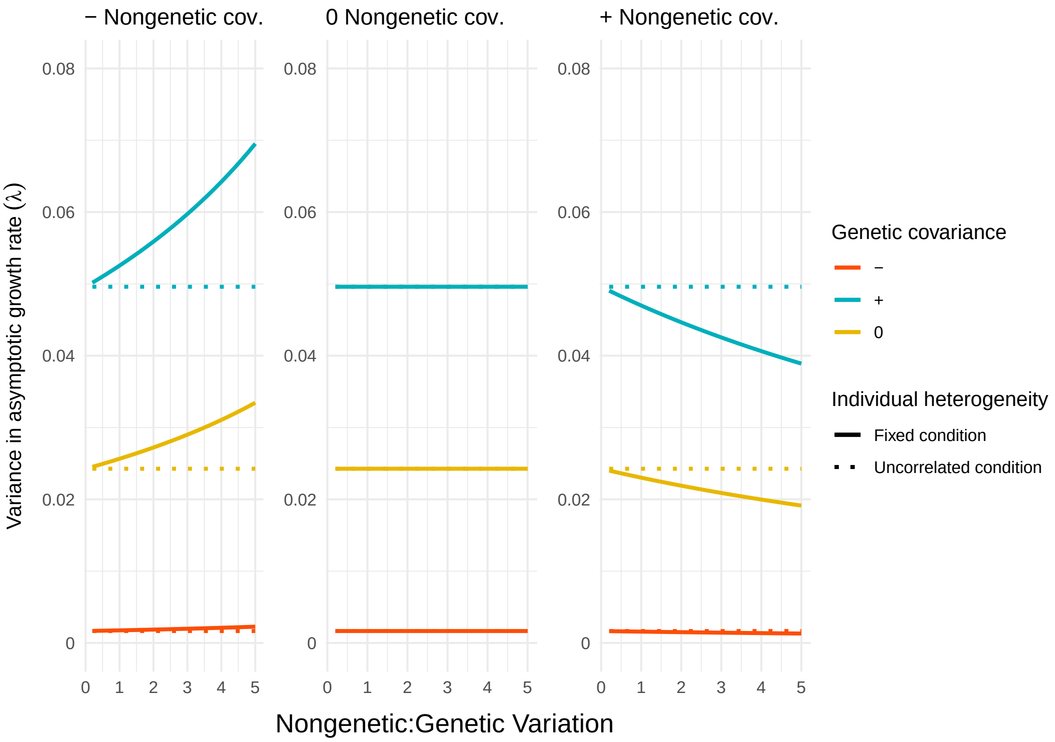


**Figure S1**. Impact of genetic and nongenetic heterogeneity on the variance in asymptotic growth rates ($\lambda$) across 1000 randomly drawn genotypes when there is early recruitment to the breeding population. The variance in asymptotic growth rates is a function of the relative strength of nongenetic variation (x-axes), the sign of the genetic and nongenetic covariance (line colours and panels, respectively), and the type of individual heterogeneity (line types). Comparing these results to fig. 2 in the main text reveals that changes in evolvability occur because both the mean and variance in asymptotic growth rates are influenced by the type and strength of individual heterogeneity when there is early recruitment. For example, evolvability declines with the relative strength of nongenetic variation when there is fixed condition and positive nongenetic covariances (right panel, fig. 2B), both because of an increase in the mean asymptotic growth rate (right panel, fig. 2A) and because of a decrease in the variance in asymptotic growth rates (right panel, above).

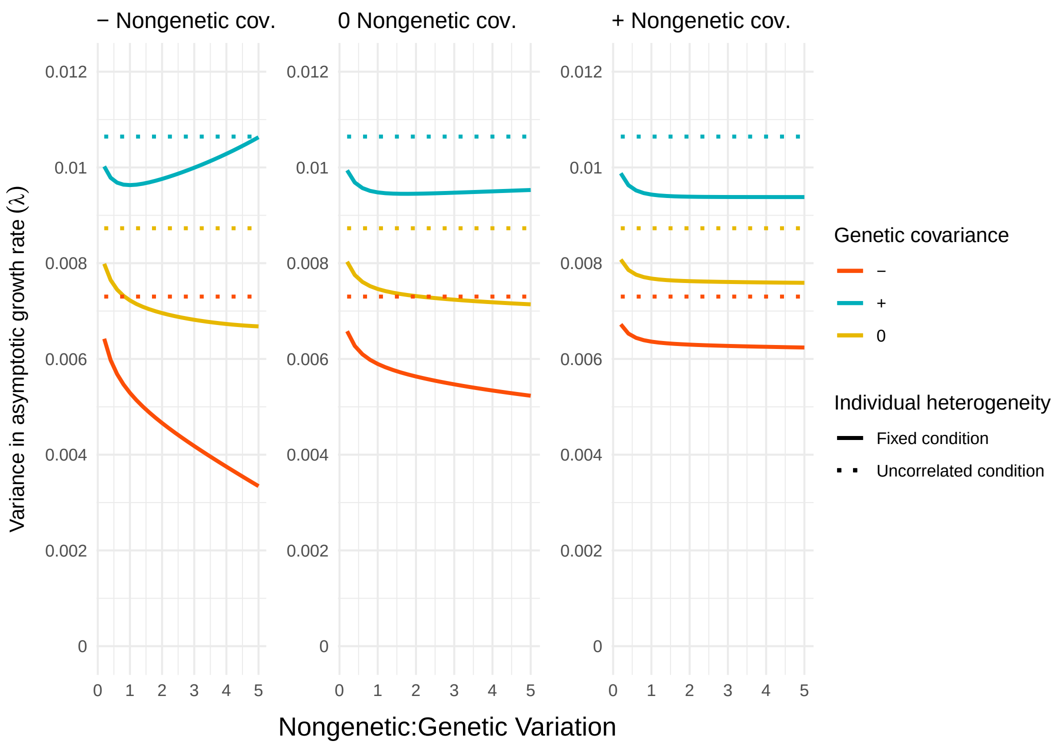


**Figure S2**. Impact of genetic and nongenetic heterogeneity on the variance in asymptotic growth rates ($\lambda$) across 1000 randomly drawn genotypes when there is late recruitment to the breeding population. The variance in asymptotic growth rates is a function of the relative strength of nongenetic variation (x-axes), the sign of the genetic and nongenetic covariance (line colours and panels, respectively), and the type of individual heterogeneity (line types). Comparing these results to fig. 3 in the main text reveals that evolvability decreases with the relative strength of nongenetic variation both because of an increase in the mean asymptotic growth rate (fig. 3A) and because of a general decrease in the variance in asymptotic growth rates (above). Note that evolvability is more strongly influenced by the mean asymptotic growth rate when there is late recruitment than when recruitment is earlier (e.g., fig. 2), causing evolvability to decline even when the variance in asymptotic growth rates increases (negative nongenetic covariance (left panel) with positive genetic covariance (blue line) above) or plateaus.

R Core Team. 2023. R: a language and environment for statistical computing. R Foundation for Statistical Computing, Vienna.
